# From breast cancer cell homing to the onset of early bone metastasis: dynamic bone (re)modeling as a driver of metastasis

**DOI:** 10.1101/2023.01.24.525352

**Authors:** Sarah A. E. Young, Anna-Dorothea Heller, Daniela S. Garske, Maximilian Rummler, Victoria Qian, Agnes Ellinghaus, Georg N. Duda, Bettina M. Willie, Anika Grüneboom, Amaia Cipitria

**Author notes:** Corresponding author Max Planck Institute of Colloids and Interfaces, Am Mühlenberg 1, 14476 Potsdam.

## Abstract

Breast cancer often metastasizes to bone causing osteolytic lesions. Structural and biophysical changes are rarely studied, yet are hypothesized to influence metastatic progression. Here, we developed a mouse model of early bone metastasis and multimodal 3D imaging to quantify cancer cell homing, dynamic bone (re)modeling and onset of bone metastasis. Using 3D light sheet fluorescence microscopy, we show eGFP^+^ cancer cells and small clusters in 3D (intact) bones. We detect early bone lesions using time-lapse *in vivo* microCT and reveal altered bone (re)modeling in absence of detectable lesions. With a new microCT image analysis tool, we detect and track the growth of early bone lesions over time. We show that cancer cells home in all bone compartments, while osteolytic lesions are only detected in the metaphysis, a region of high (re)modeling. Our study provides novel insights of dynamic bone (re)modeling as a driver during the early phase of metastasis.

## Introduction

Bone metastases often originate from breast cancer, one of the leading causes for cancer-associated deaths among women worldwide^1^. 65-80% of patients with advanced breast cancer develop bone metastasis^2, 3^. Breast cancer cells homing to bone lead to osteolytic lesions^2^ and drastically lower the prognosis of survival^4^. Metastatic cancer undergoes different stages of activity, from a dormant invisible stage that can last up to decades, to small growing metastases and up to advanced overt metastasis and skeletal morbidity^5–7^. Much work has been reported on primary tumor growth and advanced overt metastasis, but less is known about the early phases of metastasis, such as the premetastatic niche^8^, the antimetastatic niche^9^, the dormant niche^10^ or the early metastatic niche^11^.

The biophysical and structural properties of the tumor microenvironment have become a key focus of research. Advanced physical science methods hold great potential to reveal physico-chemical alterations that modulate early phases of metastasis. A recent study showed that the extracellular matrix (ECM) composed of tumor-derived collagen type III sustained tumor cell dormancy, and changes in its architecture and density restored proliferation ^12^. As another ECM protein, fibronectin fibers were found to be under lower tension in tumor invaded tissues compared to healthy tissue^13^. In the context of mineralized bone tissue, mineral nanocrystals composed of less-mature hydroxyapatite were associated with the initiation of metastasis ^11^. A follow-up study recapitulated the premetastatic niche by systemic injection of tumor conditioned media, which induced increased bone formation of altered quality^8^ and suggested potential consequences on the initial seeding of breast cancer cells. However, multimodal 3D imaging is required to shed light on the tight interlink between tissue structural changes and cancer progression during early phases of metastasis. This is particularly challenging in mineralized bone tissue, since it is less accessible than soft organs and imaging of soft bone marrow compartments within 3D intact bones requires advanced multimodal 3D imaging methods^14, 15^. The dynamics of matrix (re)modeling is another key aspect of metastatic progression^16^. Bone is a highly dynamic organ and mineralized tissue is constantly (re)modelled^17^. The differences in bone turnover between young and older animals did not affect the seeding of breast cancer cells but influenced the development of overt osteolytic bone metastasis: young mice with higher bone turnover exhibited higher numbers of overt skeletal lesions, as quantified by histology and *ex vivo* microcomputed tomography (microCT)^18^. Another study used longitudinal *in vivo* microCT to investigate the altered bone tissue dynamics due to metastasis and quantified bulk bone destruction upon direct intratibial injection of cancer cells^19^. However, the spatial resolution of the *in vivo* microCT scans limited the quantitative analysis to large bone lesions during advanced overt metastasis. In this context, we have recently described quantitatively anatomical site-specific differences in bone (re)modeling of healthy 12-week-old skeletally mature mice, with the metaphysis (in femur and tibia) being the region of highest bone (re)modeling and the tibia being more (re)modeled than the femur^20^. Furthermore, this served as baseline to detect the onset of bone pathologies, taking breast cancer metastasis as proof-of-concept^20^. Understanding the dynamics of bone (re)modeling and correlating this with the visualization and quantification of homing of breast cancer cells, will shed light on the role of bone tissue dynamics on breast cancer cell homing and/or on initiation of osteolytic lesions.

We hypothesized that the dynamic microenvironment in the bone drives the progression of early bone metastasis. Therefore, we established an experimental breast cancer bone metastasis mouse model to study the early stages of metastasis, from breast cancer cell homing to the onset of early bone lesions, both *in vivo* and *ex vivo*. Skeletally mature 12-week-old female BALB/c nude mice were injected intracardially with eGFP^+^ bone-tropic breast cancer cells, to recapitulate the stage of disseminated cancer cells in circulation. We visualized and quantified the homing of breast cancer cells and small clusters to the bone marrow with advanced light-sheet fluorescence microscopy (LSFM) using 3D (intact) long bones. LSFM revealed breast cancer cells and small clusters disseminated to all bones and different compartments. Dynamic bone (re)modeling was investigated by *in vivo* microCT-based time-lapse morphometry and revealed altered bone (re)modeling in absence of detectable bone lesions. The method was further adapted and we developed a new image analysis tool to detect and track the growth of early bone lesions over time. Osteolytic lesions occurred in characteristics locations of high bone (re)modeling activity in the metaphysis, in both cortical and trabecular bone; no lesion was detected in the epiphysis or diaphysis. In addition, we used advanced *ex vivo* multimodal methods to characterize the structural changes caused by early lesions. Altogether, the results suggest that dynamic bone (re)modeling is a driver of metastasis, as breast cancer cells reach every bone compartment, yet lesions only occur in the dynamic regions like the metaphysis. Our study provides novel insights of dynamic bone (re)modeling in metastatic initiation and progression.

## Results

### Breast cancer cells and small clusters homing in the bone marrow visualized by 3D light-sheet fluorescence microscopy

LSFM was used to study the homing of breast cancer cells and small cell clusters in 3D (intact) tibiae. Optical clearing makes the bone transparent to LSFM and allows 3D visualization of fluorescently labeled (eGFP^+^) breast cancer cells in the bone marrow and in the periosteum. The (virtual) 2D section overview image in Fig. 1A shows breast cancer cells and small cell clusters infiltrating the entire bone marrow of the tibia (Fig. 1A.1-2) (Supplementary Movie 1). In addition, larger and denser clusters of cancer cells were found in the periosteal region (Fig. 1A.3). Single cells and small clusters of only a few cells were found in between muscle tissue and all over the periosteal bone surface (Fig. 1A.4). The quantitative analysis of the size and spatial distribution of around 5,500 cells and cell clusters in the bone marrow of five different bones (Fig. 1F) shows that most breast cancer cells are in close proximity to the cortical bone surface (Supplementary Movie 2), with a mean distance of around 50 µm (Fig. 1C) and a mean diameter ranging between 31-36 µm (Fig. 1D). Periosteal cell clusters on the other hand can be much larger with volumes up to 0.1 mm^3^ (Fig. 1E).

**Fig. 1.**
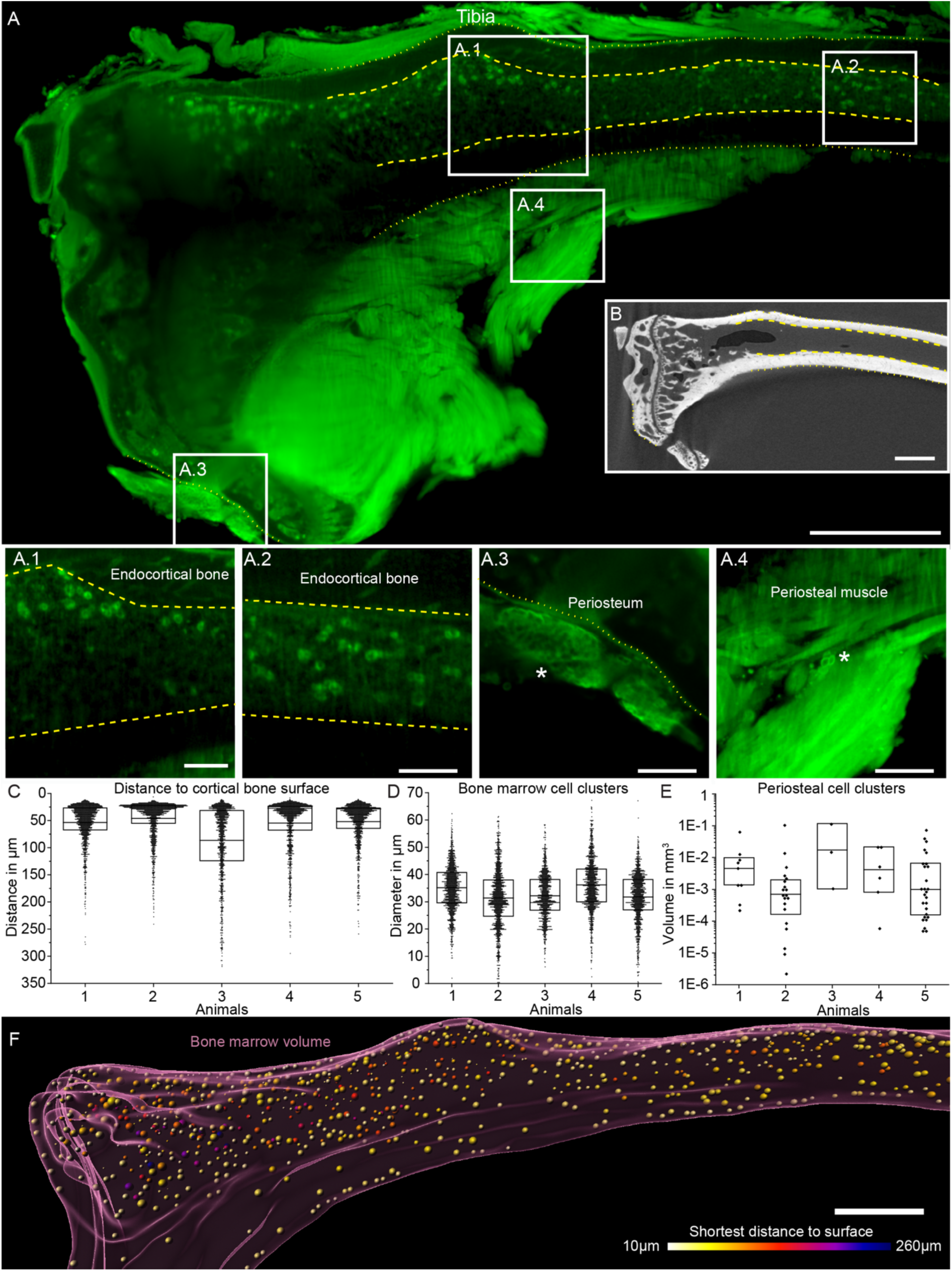
Light-sheet fluorescence microscopy (LSFM) and 3D quantification of the size and spatial distribution of breast cancer cells and small clusters homing in the bone marrow. LSFM images of one (virtual) section are depicted for (A) one entire bone, with enlargements of small cell clusters in the (A.1) bone marrow upper diaphysis with the endocortical bone surface delimited with a dashed yellow line, (A.2) bone marrow lower diaphysis, (A.3) periosteal site with a large dense cluster (asterisk) and (A.4) in between muscle tissue with a small cluster (asterisk). (B) MicroCT image of the same (virtual) bone section as shown in A, with highlighted endocortical and periosteal bone surfaces. Quantification of (C) the distance to the cortical bone surface and (D) the size distribution of breast cancer cells and small clusters. (E) Quantification of the volume of larger breast cancer cell clusters on the periosteum. (F) Graphical 3D visualization of the distance of breast cancer cells and small clusters to the endocortical bone surface, with the color code indicating the distance. (Asterisks in A.3 and A.4 indicate the presence of cancer cells and scale bars correspond to 500 µm (A and B) and 200 µm (A.1-A.4) respectively. The box plots show the mean (C and D) or median (E) and first and third quartile. n = 5 bones from five tumor animals, representative images from one animal with a trabecular bone lesion).

Further descriptive analysis of the homing of breast cancer cells and small clusters revealed that these are detected in all bone compartments like the diaphysis, metaphysis and epiphysis, as well as in different types of bone such as tibia and fibula, as shown in Fig. 2.

**Fig. 2.**
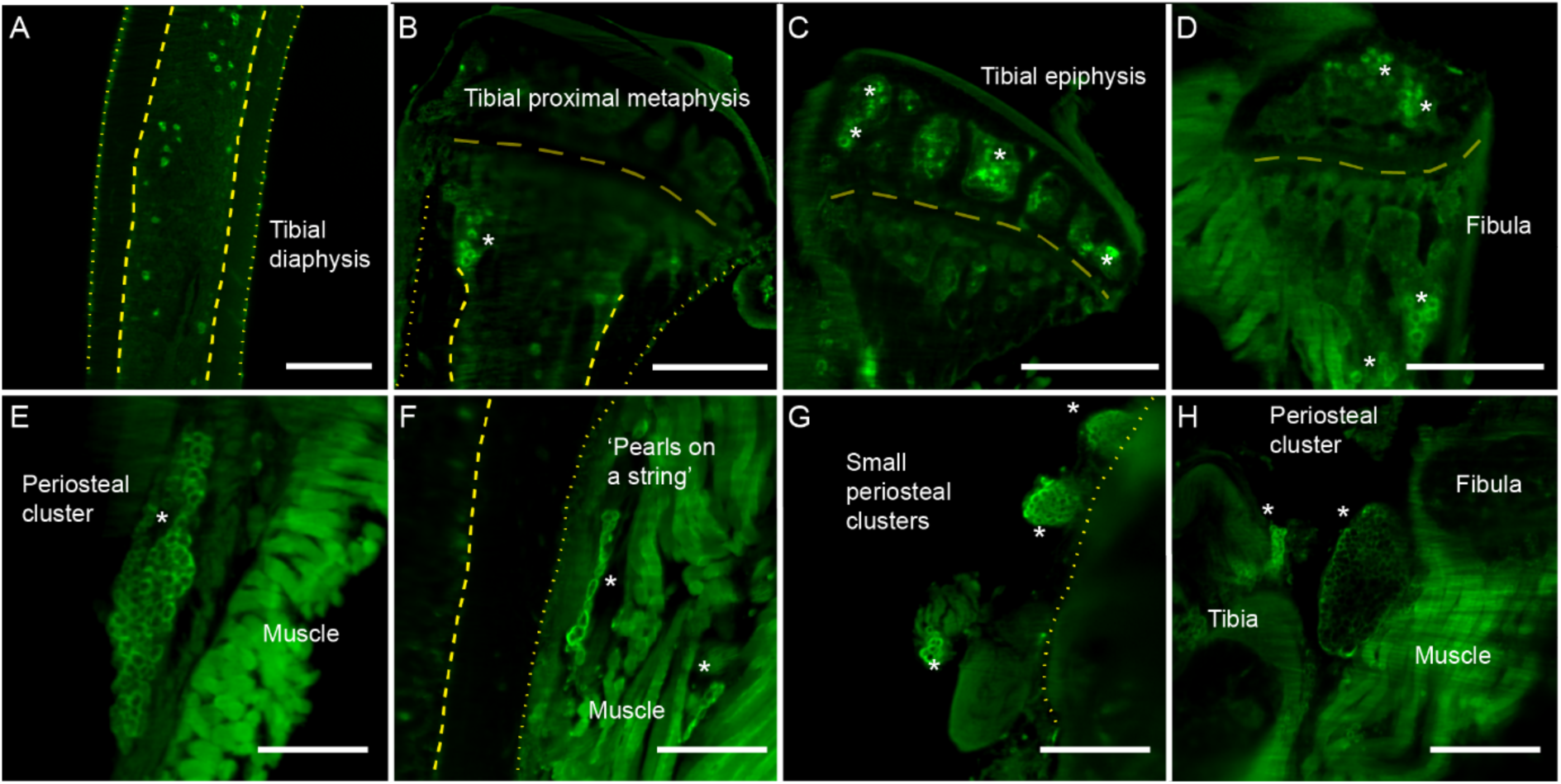
LSFM visualization of breast cancer cells and small clusters in various compartments of skeletal tissue. Images showing cancer cells in the (**A**) tibial diaphysis, (**B**) tibial proximal metaphysis, (**C**) tibial epiphysis and (**D**) fibula, as well as in the periosteal tissue as (**E**) a large cluster, (**F**) like ’pearls on a string’, (**G**) as small clusters and (**H**) as a large cluster between tibia and fibula. (Scale bars correspond to 500 µm (A-D) and 250 µm (E-F), cancer cells and patches are marked with an asterisk; periosteal and endocortical bone, as well as the growth plate are delimited with dashed yellow lines. n = 5 bones from five tumor animals).

### Detection and tracking of the growth of early bone metastatic lesions over time

To detect the location and track the growth of early metastatic lesions over time in cortical bone, a new image analysis tool was developed, using *in vivo* microCT-based time-lapse morphometry. Dynamic bone (re)modeling was investigated by overlapping a reference scan (4 days prior to cancer cell injection) with later time points. The resulting visualization of constant bone (yellow), newly mineralized bone (blue) and eroded bone (red) can be quantified in volume, surface and apposition/resorption rate^20, 21^.

Our new image analysis tool has the ability to differentiate between physiological and pathological erosion patches. The method is based on empirically determined threshold parameters to define a 3D neighborhood by summing up eroded voxels as shown in Fig. 3A. Briefly, a first noise filtering is performed and only eroded bone patches are considered, which fulfill the criterion that 70% of the neighborhood (respectively 3 voxels in positive and negative x-, y- and z-direction) consists of erosion voxels (Fig. 3B). Next, to distinguish physiological vs. pathological erosion patches, we search for the pathological eroded bone patch core, which fulfills the stricter criterion that 92% of a larger neighborhood (respectively 9 voxels in positive and negative x-, y-direction, 4 voxels in positive and negative z-direction) consists of eroded voxels (Fig. 3C). Finally, the volume of the pathological eroded bone patch is determined by the noise-filtered eroded bone patch including the patch core (Fig. 3D). The threshold settings were obtained and validated with datasets from PBS-injected control animals with physiological bone (re)modeling, confirming that no pathological eroded bone patches were detected.

**Fig. 3.**
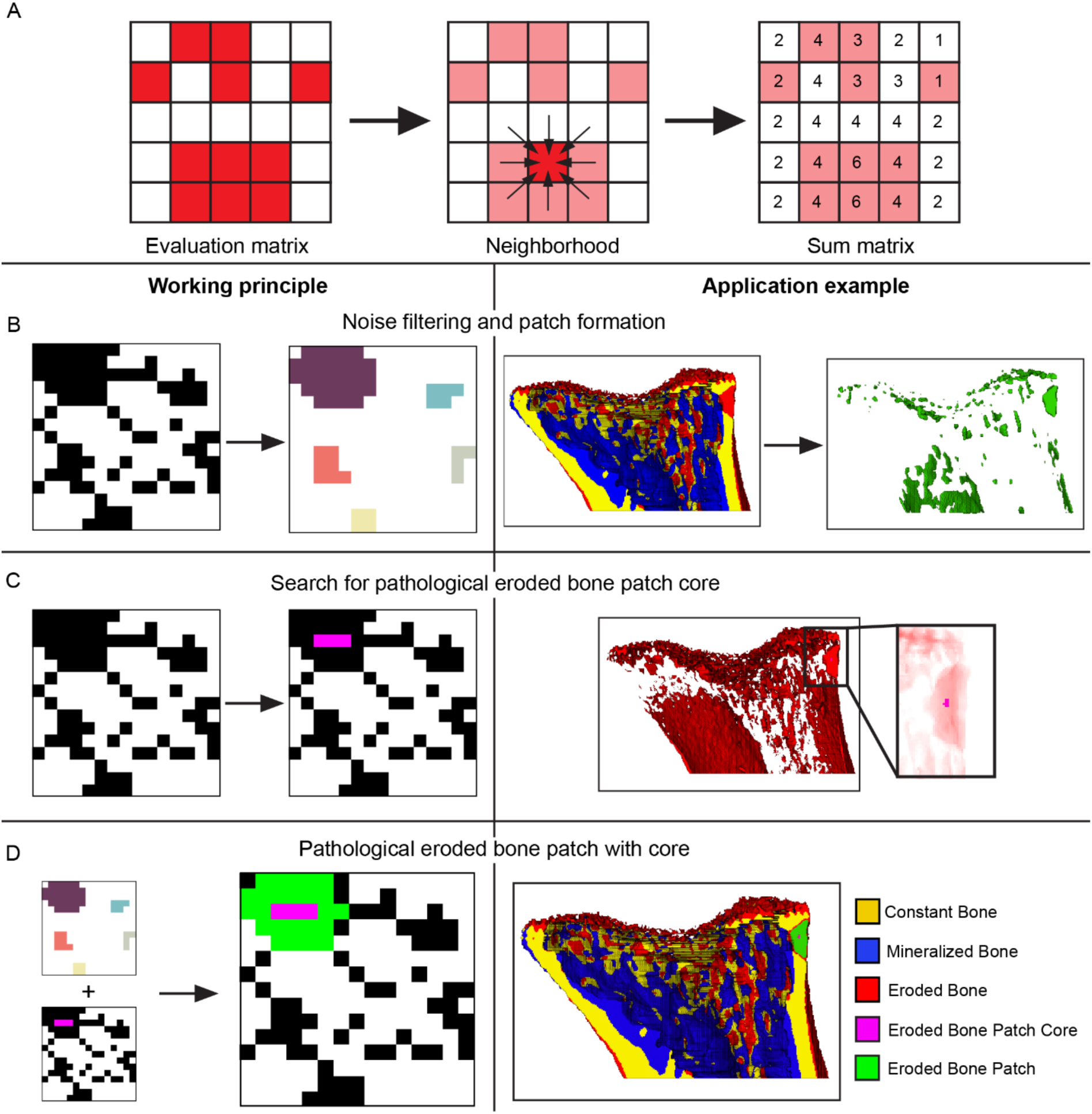
Eroded bone patch image analysis tool to detect and track the growth of early bone metastatic lesions over time based on in vivo microCT time-lapse morphometry. (**A**) Schematic showing how the neighborhood is determined, starting at the evaluation matrix, with a 2D simplified 3×3 neighborhood for one voxel and the resulting sum matrix for all voxels. Working principle (left) and application example (right) of the eroded bone patch analysis starting with (**B**) filtering of noise to form separate patches, (**C**) search for a pathological eroded bone patch core and (**D**) determining the volume of the pathological eroded bone patch by matching the filtered patch with the core. Constant bone is shown in yellow, mineralized bone in blue, eroded bone in red, eroded bone patch in green and eroded bone patch core in pink.

Osteolytic erosion patches were found in the cortical bone of six animals, with three in the tibia and three in the femur as shown in Fig. 4 and one exemplary osteolytic lesion in Supplementary Movie 3 and Movie 4. Five of these lesions started on the periosteal bone surface, suggesting an onset in that region (Fig. 4A, B, D) and one in the endocortical bone (Fig. 4C). All lesions were detected as early as day 17, which is the first time point that was measured after the reference scan (day 0, four days prior to cancer cell injection). The patch size at first detection varied between 0.005 – 0.025 mm^3^, with equal distribution of smaller and bigger lesions in both femur and tibia. Lesions were tracked for up to 27 days (animal sacrifice due to humane endpoint criteria). In several animals the osteolytic eroded bone patch volume increased, with the clearest example being animal 1 starting at 0.01 mm^3^ and reaching a final lesion volume of nearly 0.1 mm^3^ (Fig. 4E). The eroded bone patch analysis enables the spatio-temporal visualization and quantification of early osteolytic lesions, which provides fundamental understanding of the onset of bone metastasis.

**Fig 4.**
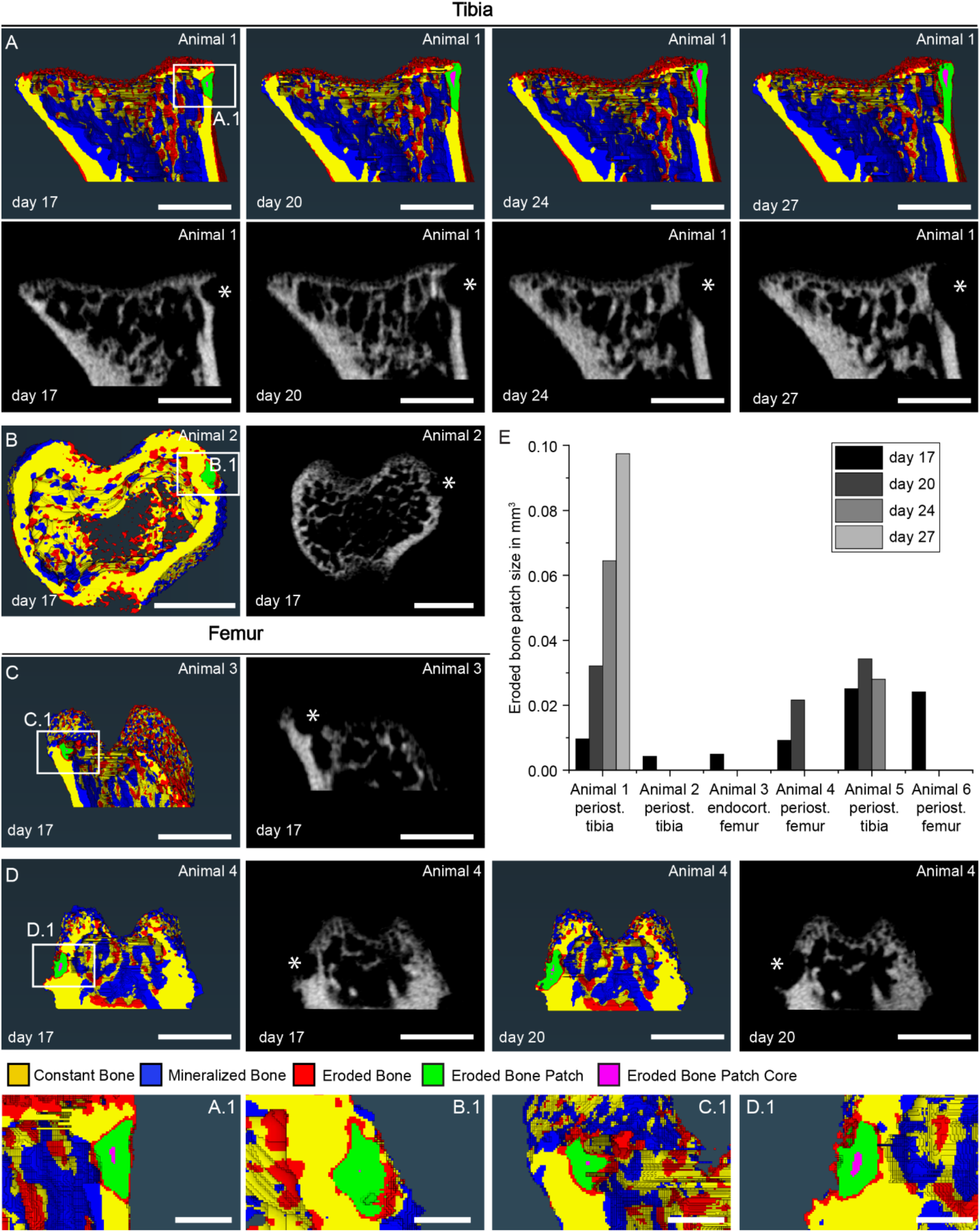
Spatio-temporal analysis of the location and growth of early metastatic lesions in cortical bone using the new eroded bone patch image analysis tool. Results of bone (re)modeling and detection of pathological erosion-patches in the tibia of (**A**) animal 1 at day 17 (enlargement in (**A.1**)), day 20, day 24 and day 27, and (**B**) animal 2 at day 17 (enlargement in (**B.1**)). Analogous analysis for the femur of (**C**) animal 3 at day 17 (enlargement in (**C.1**)) and (**D**) animal 4 at day 17 (enlargement in **D.1**) and day 20. (**E**) Quantitative results of the eroded bone patch analysis per animal up to 27 days. (Bone (re)modeling visualization of constant bone in yellow, mineralized bone in blue, eroded bone in red, eroded bone patches in green, eroded bone patch core in pink and respective microCT grey scale images. Scale bars correspond to 1 mm, scale bar for enlargements 0.2 mm. The lesions are indicated with an asterisk in the grey scale images. n = 6 bones from six tumor animals).

### Metastatic lesions in trabecular bone initiate on the mineralized surface adjacent to the growth plate

Metastatic osteolytic lesions in trabecular bone were visually detected in the primary spongiosa, right on the mineralized surface adjacent to the growth plate, mainly in the tibia (Fig. 5A-C) but also in the femur (Fig. 5D). For all lesions, a top view and enlargements of a cross section are shown, both as bone (re)modeling analysis data and the respective grey scale image. The enlarged images (Fig. 5A.1-D.1) confirm that the growth plate is eroded in a larger area, spanning both along the growth plate and towards the secondary spongiosa. Although all lesions are still at a very early stage, Fig. 5B.1 clearly shows the smallest lesion with a diameter along the growth plate of under 200 µm. Two of the lesions are right between the condyles (Fig. 5A and D), the others do not show any distinct pattern.

**Fig. 5.**
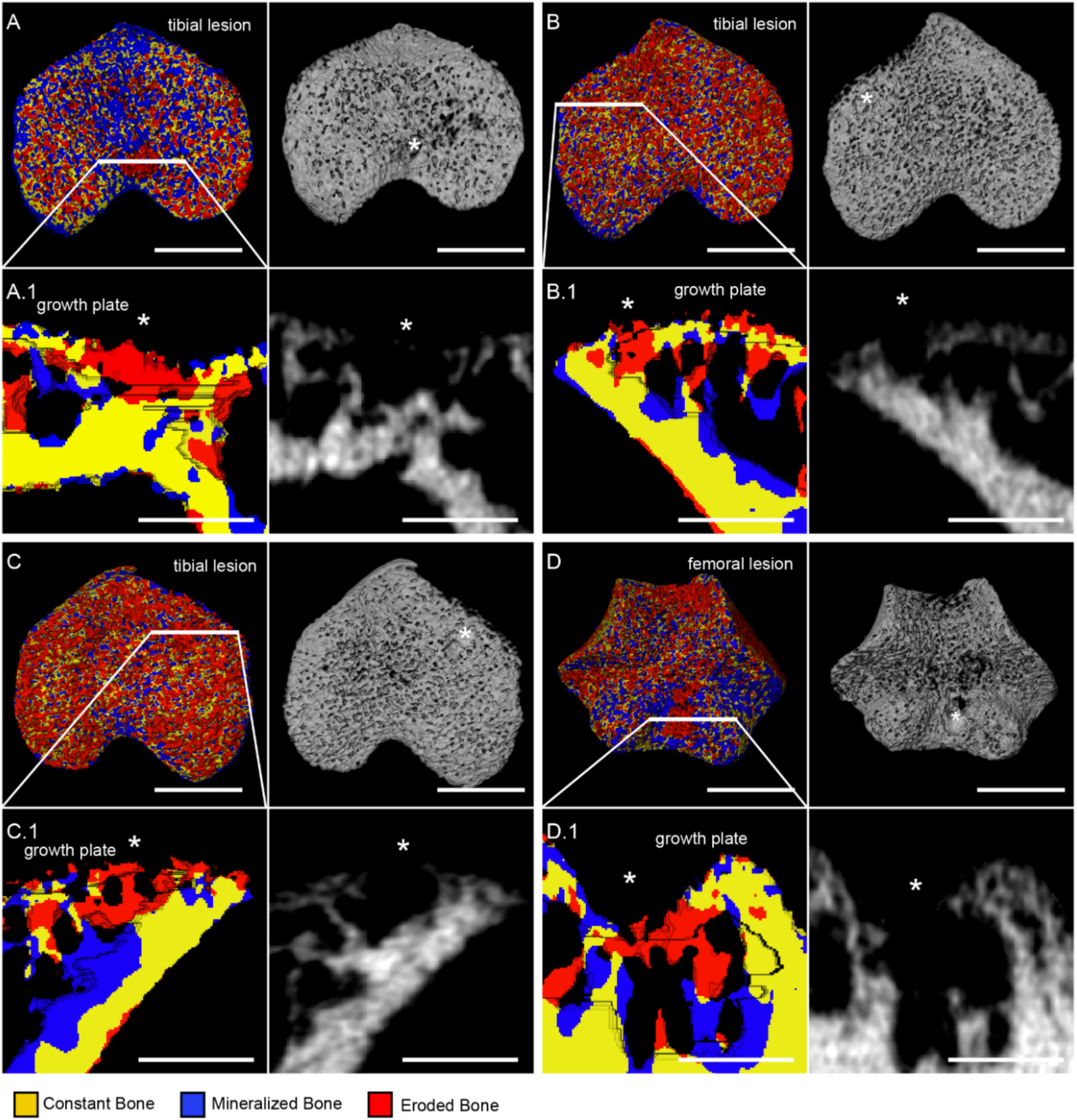
Early metastatic lesions in trabecular bone initiation at the mineralized growth plate in the primary spongiosa as shown by microCT-based time-lapse morphometry. Results of bone (re)modeling with constant bone in yellow, mineralized bone in blue, eroded bone in red, as well as grey scale images for all bones at day 17. (**A**) (Re)modeling visualization top view, with respective grey scale image and (**A.1**) enlargement of the cross-section of the osteolytic lesion of a tibia (**B**) Analogous visualization and (**B.1**) enlargement of the cross-section of a tibia. (**C**) Analogous visualization and (**C.1**) enlargement of the cross-section of a tibia. (**D**) Analogous visualization and (**D.1**) enlargement of the cross-section of a femur. (Scale bars correspond to 1 mm and 0.5 mm for enlargements. The lesions are indicated with an asterisk. n = 4 bones from two tumor animals).

To summarize, we were able to identify the onset of metastatic bone lesions visualized with microCT-based time-lapse morphometry, on both cortical and trabecular bone. Based on the location where they initiate, we identify: (i) fully cortical lesions that start on the periosteal side and do not reach the trabecular bone even after growing for 10 days (Fig. 4A); (ii) mixed lesions where both the endocortical bone (Fig. 4C) and adjacent trabecular bone are resorbed (Fig. 5D); (iii) fully trabecular lesions located directly at the mineralized growth plate in the metaphysis (Fig. 5).

### Structural microenvironment of early metastatic bone lesions visualized with multiscale correlative tissue characterization

The early metastatic bone lesions detected with microCT-based time-lapse morphometry were then subjected to multiscale correlative tissue characterization to evaluate the structural microenvironment. Cortical and trabecular bone osteolytic lesions in femur (Fig. 6A-I) and tibia (Fig. 6J-R) were analyzed and compared with intact regions within the same bone.

**Fig. 6.**
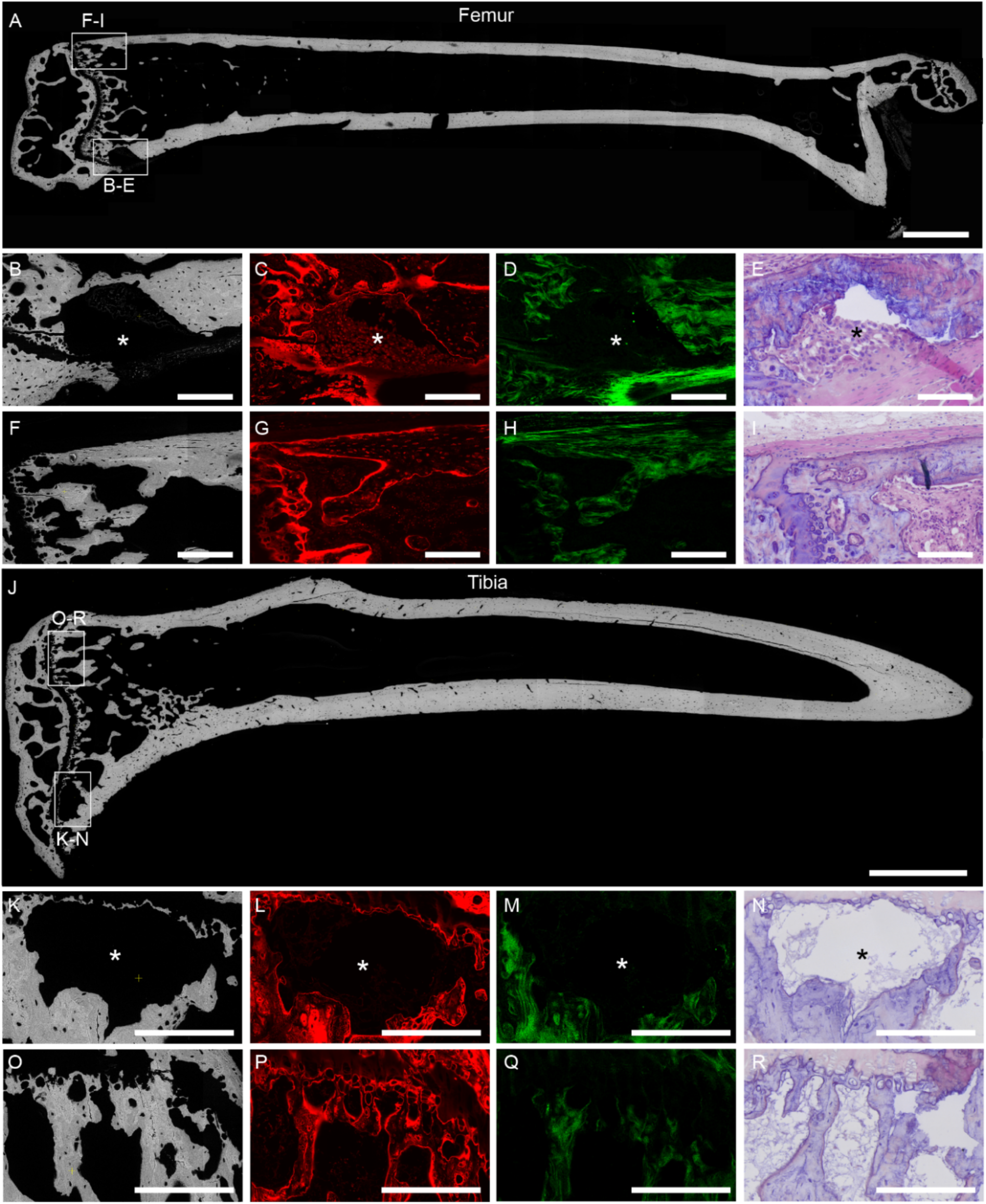
Multiscale correlative tissue characterization of early metastatic lesions in cortical and trabecular bone and comparison with intact bone tissue in the same sample. (**A**) Overview BSE microscopy image of a femur with (**B**) enlargement of a cortical lesion, as well as (**C**) CLSM of rhodamine-stained tissue, (**D**) SHG imaging of collagen fibers and (**E**) H&E histology. (**F**) BSE microscopy enlargement of an intact bone region, as well as (**G**) CLSM of rhodamine-stained tissue, (**H**) SHG imaging of collagen fibers and (**I**) H&E histology. (**J**) Overview BSE microscopy image of a tibia with (**K**) enlargement of a trabecular lesion, as well as (**L**) CLSM (rhodamine staining), (**M**) SHG imaging of collagen fibers and (**N**) H&E histology. (**O**) BSE microscopy enlargement of a healthy trabecular region, as well as (**P**) CLSM (rhodamine staining), (**Q**) SHG imaging of collagen fibers and (**R**) H&E histology. (Asterisks indicate the presence of a lesion, scale bars correspond to 1 mm and 200 µm in the enlargements. n = 5 bones from five tumor animals containing an osteolytic lesion were analyzed, representative images from two animals with a cortical lesion in the femur and a trabecular lesion in the tibia are shown).

An overview image of the femur using backscattered electron (BSE) microscopy is shown in Fig. 6A, with the mineralized tissue of the bone being intact except for the cortical lesion in the posterior metaphysis (enlargement in Fig. 6B). The cortical bone is resorbed across nearly the entire cortical thickness, with only a thin layer remaining on the endocortical side. In the progression of such early osteolytic lesions, the matrix containing the lacuno-canalicular network is resorbed (Fig. 6C). The collagen analysis with second-harmonic generation (SHG) imaging also shows that all fibrillar collagen type I fibers that were part of the cortical bone have been resorbed (Fig. 6D). Interestingly, the thick collagen layer in the periosteum is still intact (Fig. 6D). The lesion region is now filled with a new soft tissue with high density of cells (Fig. 6C, E).

Similarly, an overview image of the tibia using BSE microscopy is shown in Fig. 6J, with the mineralized tissue of the bone being intact except for the trabecular lesion on the anterior growth plate (enlargement in Fig. 6K). The mineralized part of the growth plate is visibly thinned out in that region and the surrounding trabeculae show a rough surface, indicating an ongoing erosion of the tissue. In analogy to the cortical lesion, the matrix containing the lacuno-canalicular network (Fig. 6L) and fibrillar collagen type I fibers (Fig. 6M) are fully resorbed, while the growth plate on the posterior side (Fig. 6O-R) is fully intact.

In summary, in both early cortical and trabecular lesions, the mineralized tissue, including embedded lacunae and canaliculi, as well as collagen fibers within mineralized tissue are all fully resorbed and replaced by a new soft tissue with high density of cells.

### Breast cancer cells and small clusters homing in the bone marrow have a tendency for low proliferation

The proliferation state of the breast cancer cells and small clusters in the bone marrow was analyzed using the proliferation marker Ki67. Sections from three bones harvested from three tumor animals, exhibiting an osteolytic lesion in at least one bone, were imaged. In total 364 breast cancer cells or small clusters were detected inside the bone marrow (Fig. 7A with enlargements in A.1, and A.2) and analyzed for the proliferation status (Fig. 7B). A cluster of cells was counted as Ki67^+^ as soon as one cell in this cluster was positive. High resolution examples of Ki67^+^ (Fig. 7A.1) and Ki67^-^ (Fig. 7A.2) are shown and single channel views are included in Fig. S1. We show that half of the single cells and small clusters are Ki67^-^, while those considered as Ki67^+^ typically entail only one Ki67^+^ cell. We conclude that at early stages of breast cancer homing to the bone marrow, single cells and small clusters have a tendency for low proliferation.

**Fig. 7.**
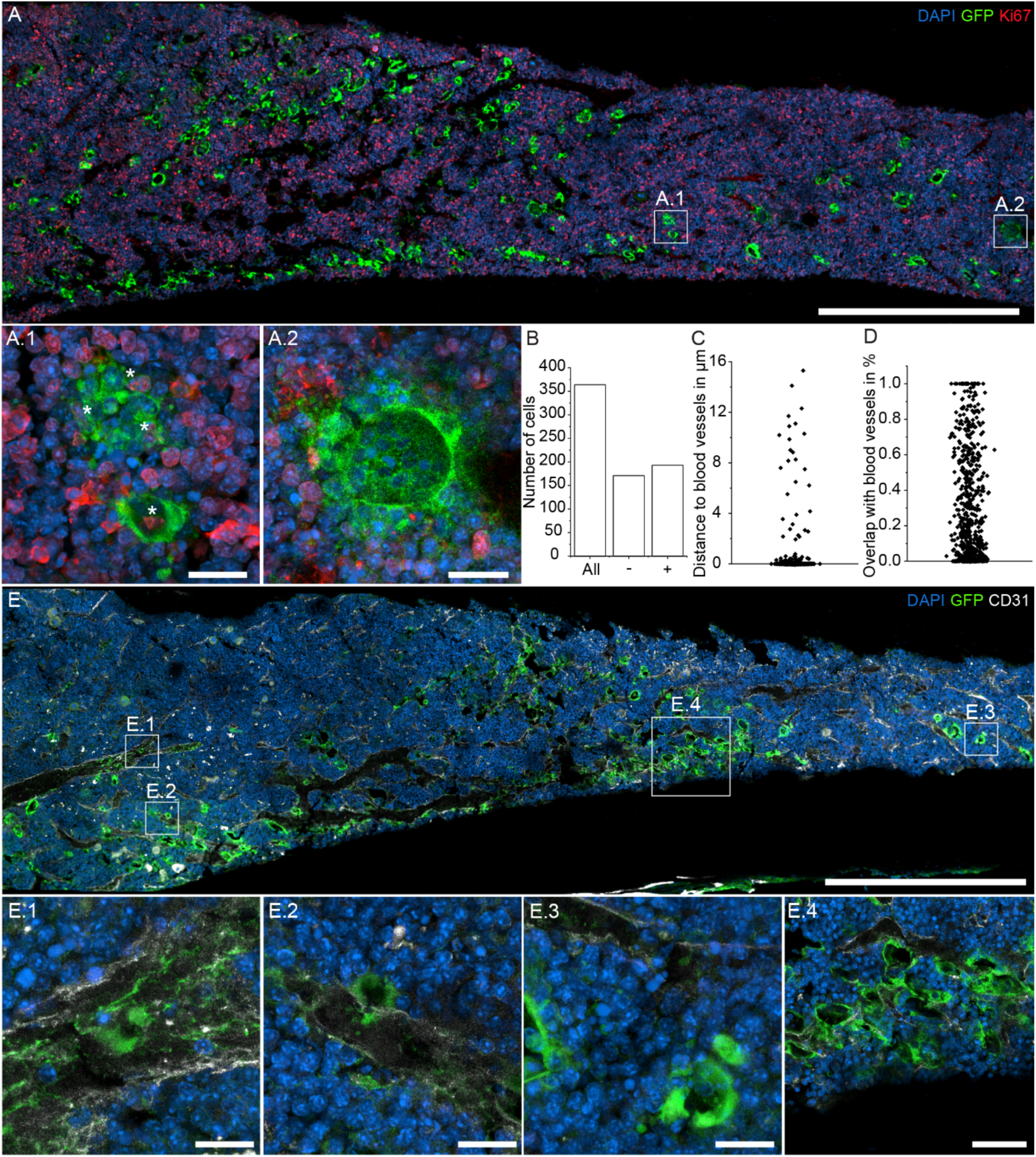
Breast cancer cells and small clusters are low in proliferation and found in close proximity and inside of blood vessels. **(A)** CLSM imaging of Ki67 proliferation marker with enlargement (**A.1**) showing a Ki67 positive breast cancer cell cluster and (**A.2**) showing a Ki67 negative breast cancer cell clusters. (**B**) Quantification of the Ki67 signal for all n = 364 analyzed single cells and small clusters. Quantification of breast cancer cell and small cluster (**C**) distance to blood vessels and (**D**) degree of overlap with blood vessels. (**E**) CLSM imaging of CD31 with enlargements showing breast cancer cells and small clusters (**E.1**) inside a blood vessel, (**E.2**) in direct contact with a blood vessel, (**E.3**) with a larger distance from blood vessels and (**E.4**) close to the endosteal bone surface. (Scale bars correspond to 500 μm and 20 μm for the enlargements, except for E.4 with 50 µm. DAPI staining of the nucleus in blue, eGFP of the cancer cells in green, Ki67 in red and CD31 in white. n = 3 bones (Ki67) from three tumor animals and n = 4 bones (CD31) from four tumor animals with osteolytic lesions in other long bones).

### Breast cancer cells and small clusters are found in close proximity to and inside of blood vessels

The distance of individual breast cancer cells and small clusters to blood vessels in the bone marrow stained with the CD31 endothelial cell marker was analyzed with the software Imaris (Fig. 7E). Sections from four bones harvested from four tumor animals, with an osteolytic lesion in at least one hind limb were analyzed. Most cancer cells can be found in direct contact with blood vessels and only few at further distance, with a maximum distance at 16 µm (Fig. 7C). Breast cancer cells are overlapping with blood vessel to a different extent as quantified in Fig. 7D, from 100% overlap (Fig. 7E.1), to partial overlap (Fig. 7E.2) or no overlap (Fig. 7E.3). In addition, cancer cells are found close to the endocortical bone surface, which is also a highly vascularized region (Fig. 7E.4).

### Homing of breast cancer cells in the bone marrow enhances new mineralization in the absence of detectable osteolytic lesions

To study the effect of breast cancer cell homing in the dynamics of bone (re)modeling, we investigated tumor animals without detectable osteolytic lesions (n = 5) up to 38 days and compared with PBS-injected control animals (n = 7). The trabecular bone (re)modeling in the tibia proximal metaphysis is shown in Fig. 8A-H. An exemplary image of a tumor animal is depicted in Fig. 8A and the PBS-injected control animal in Fig. 8B. The corresponding mineralized volume (MV/BV), surface (MS/BS), with the soft tissue interface (MS_ST_/BS) and constant bone interfaces (MS_CB_/BS)^20^ and mineral apposition rate (MAR) are shown in Fig. 8C-E, while the eroded volume (EV/BV), surface (ES/BS), with the soft tissue interface (ES_ST_/BS) and constant bone interfaces (ES_CB_/BS)^20^ and mineral resorption rate (MRR) in Fig. 8F-H. The tumor animal shows more newly mineralized trabecular bone and less eroded bone (Fig. 8A), compared to the PBS-injected control animal.

**Fig. 8.**
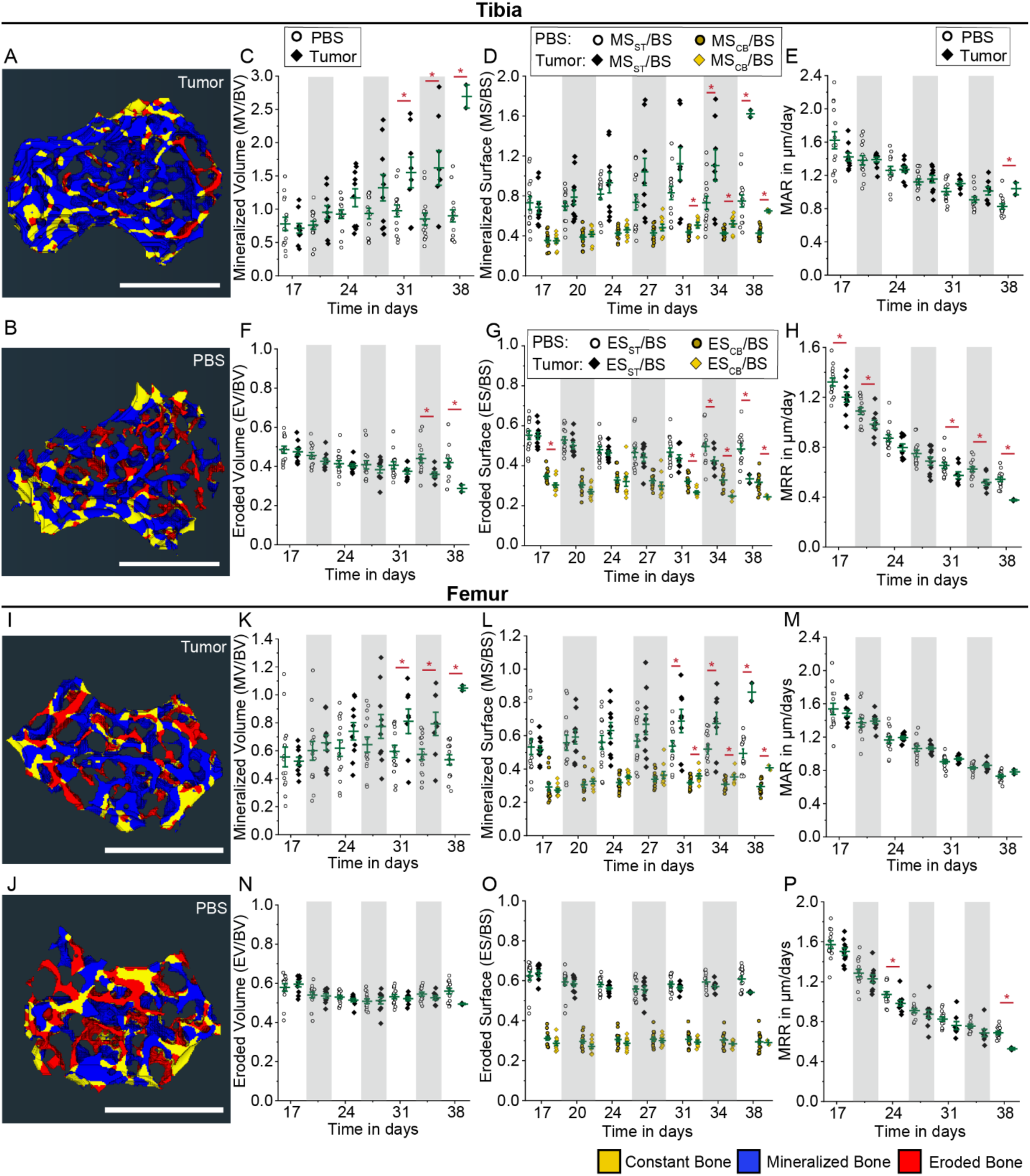
Altered trabecular bone (re)modeling in absence of detectable bone lesions. Results in trabecular bone of tibial proximal metaphysis are depicted for (**A**) tumor cell injected animals and (**B**) PBS-injected control animals, with newly mineralized bone in blue, eroded bone in red and constant bone in yellow. Results are shown for mineralization with (**C**) MV/BV, (**D**) MS_ST_/BS as well as MS_CB_/BS and (**E**) MAR, as well as erosion with (**F**) EV/BV, (**G**) ES_ST_/BS as well as ES_CB_/BS and (**H**) MRR. Analogous analysis for trabecular bone in the femoral distal metaphysis depicted for (**I**) tumor cell injected animals and (**J**) PBS-injected control animals. Results are shown for mineralization with (**K**) MV/BV, (**L**) MS_ST_/BS as well as MS_CB_/BS and (**M**) MAR, as well as erosion with (**N**) EV/BV, (**O**) ES_ST_/BS as well as ES_CB_/BS and (**P**) MRR. (All plots show mean and standard deviation, with statistically significant values presented as * p ≥ 0.05, a two-sample t-tests was used. Exemplary images were taken from registration data at day 34, scale bars correspond to 1 mm. PBS control group: n = 14 bones from seven animals, Tumor group: n = 10 bones from five animals, for tibia and femur respectively.)

Higher MV/BV is observed in the tumor animals at all time points (except day 17), with values reaching between up to 1.5 – 2.5 for tumor animals, while it stays below 1 for PBS-injected control animals, and showing significant differences after day 31 (Fig. 8C). Similar results are observed for MS/BS (Fig. 8D), while the MAR is only significantly elevated at day 38 (Fig 8E). Lower erosion in the tumor animals can be observed with EV/BV (Fig 8F) and ES/BS (Fig. 8G), showing significant differences after day 34, and MRR (Fig. 8H) at day 17, 20, 31, 34 and 38. Analogous analysis is performed in the femur distal metaphysis (Fig. 8I-P). Trabecular bone volume, surface and apposition/resorption rate are shown in Fig. 8K-P, with exemplary images for the tumor animals in Fig. 8I and PBS-injected control animals in Fig. 8J. Similar trends as in the tibia are found in the femur, although the femur shows overall less (re)modeling. Similar trends are also found in the cortical bone (Fig. S2), with overall less (re)modeling.

## Discussion

In this study, we aimed to understand the role of biophysical and structural properties of the bone microenvironment and its dynamics on the early phases of breast cancer bone metastasis. To do so, we developed an early bone metastasis mouse model using skeletally mature 12-week-old female BALB/c nude mice. We visualized and quantified the spatial distribution of disseminated breast cancer cells in (intact) bone by 3D light sheet fluorescence microscopy. Further, we quantified dynamic bone structural changes using *in vivo* microCT-based time-lapse morphometry and developed an image analysis tool to detect and track early osteolytic lesions. Combining multimodal advanced 3D imaging, we revealed that breast cancer cells disseminate in all bone compartments, yet bone osteolytic lesions initiate in regions of high dynamic bone (re)modeling. Our study provides novel insights of the interplay between dynamic bone structural changes and metastasis initiation and progression.

Using tissue clearing and 3D light sheet fluorescence microscopy, we visualize for the first-time eGFP^+^ bone-tropic breast cancer cells and small clusters in (intact) 3D mineralized bones (Fig. 1) and show that they home to all bone compartments, including the epiphysis, metaphysis and diaphysis (Fig. 2). Quantitative analysis of their size and spatial distribution within the bone marrow revealed that the breast cancer cells and small clusters range between 10-60 μm and are in close proximity to bone surfaces. Larger clusters are found on the periosteum with a volume of up to 0.1 mm^3^. This might indicate that only small clusters or single cells can traverse capillaries and home in the bone marrow. In addition, the large clusters on the periosteum could be a sign of a more advanced stage of metastasis and are well in line with the osteolytic lesions starting at the periosteum (Fig. 5). The ability of cancer cells to form clusters has been described in the past and it was associated with a higher potential to survive and form lesions than single cells^22^.

In contrast with the finding that breast cancer cells home to all bone compartments, the localization of osteolytic lesions is highly specific in regions of high bone (re)modeling like the metaphyseal bone (Fig. 4-6), while no lesions were detected in the epiphysis or the diaphysis. Therefore, the ability for breast cancer cells to home to the bone is not sufficient as an explanation for the formation of osteolytic lesions. Cortical and trabecular bone in the metaphysis undergoes more (re)modeling than the diaphysis^21^ or the epiphysis^20^. This calls for a new time dependent and dynamic criterion for metastatic initiation and progression. This is in agreement with previous studies highlighting the importance of animal age in bone metastasis progression: young mice with higher bone turnover exhibited higher numbers of overt skeletal lesions, despite having comparable number of breast cancer cells disseminated to the bone as in older animals^18^. In our study, we used skeletally mature 12-week-old mice. Yet, even in skeletally mature mice, there are region-specific differences (metaphysis over diaphysis or epiphysis) and anatomic differences (tibia metaphysis vs. femur metaphysis) in bone (re)modeling as shown previously^20^ that could be a driver for metastasis.

To detect and track the location and growth of early metastatic lesions in cortical bone over time, we developed the new eroded bone patch image analysis tool (open code in GitHub). This allowed us to reliably find cortical osteolytic lesions as small as 0.004 mm^3^ and as early as two weeks after cancer cell injection (first time point measured). The tool was tested with PBS-injected control animals for up to six weeks post-injection, to rule-out the detection of physiological erosion patches. A similar tool was developed in the past by Campbell et al., where a distance transformation was used to detect lesions with a lowest distance of 150 μm for all planes (in x, y and z)^23^, omitting the specific characteristic of lesions to grow across the cortical thickness (in x and y, to a lesser extent in z). In addition, their tool was not tested on healthy mice that undergo major (re)modeling especially at early ages (before 12 weeks). Our eroded bone patch analysis tool has proven reliable in finding early small osteolytic lesions and, importantly, without detecting physiological eroded bone patches. We identified two different kinds of osteolytic lesions in cortical bone: on the periosteal bone surface and in the endocortical region. The majority of cortical lesions initiated at the periosteal side (n = 5 out of 6). This type of cortical lesion had not been reported in the past. The second type in the endocortical region involved erosion of both cortical and trabecular bone.

Next to cortical lesions, we analyzed early osteolytic lesions in trabecular bone. All of the lesions were visually detected two weeks after the cancer cell injection (earliest time point) and were located at the growth plate area, eroding the mineralized primary spongiosa (n = 4, three in the tibia, one in the femur). The growth plate is one of the most (re)modeled areas within the bone, and in particular in the tibia, as shown in our previous study with a quantitative analysis of spatial gradients in metaphyseal bone (re)modeling in tibia vs. femur^20^. This further supports the idea that local bone (re)modeling could be a driver for metastasis. One limitation in our study is that trabecular lesions were detected visually based on of both original microCT images and microCT-based time-lapse morphometry; the new eroded bone patch image analysis tool is only applicable to cortical bone. This probably led to an underestimation of trabecular lesions. The results we have shown here and the techniques established for cortical bone are promising for future development of additional tools to (semi)automatically detect osteolytic lesions in trabecular bone, which has a more complex geometry and undergoes more (re)modeling than cortical bone^20^.

Upon detection and 3D localization of microscopic metastatic lesions as early as two weeks after cancer cell injection, we were able to perform multiscale correlative characterization of the organic and inorganic ECM at the onset of metastatic lesions. As representative regions, we chose a cortical lesion starting at the periosteum and a trabecular lesion adjacent to the growth plate (Fig. 6). Those early lesions first identified with *in vivo* microCT-based time-lapse morphometry, were then captured on exposed 2D bone surface with controlled-angle cutting^24^ and confirmed with BSE microscopy of the exposed 2D bone surface. Previous work focusing on skeletal metastatic lesions showed animals with massive lesions, where much of the mineralized tissue had been eroded 4-6 weeks after injection and, hence, no clear analysis of the onset of the lesion could be made^11^. Combining BSE with rhodamine staining and CLSM, as well as SHG imaging, we were able to show that even in small bone metastatic lesions, the collagen fibers and the entire mineralized tissue were resorbed. As a consequence, important structures of the lacuno-canalicular network were disrupted, especially in the case of cortical lesions, spanning the entire cortical thickness. The lacuno-canalicular network plays a key role in bone mechanosensation through fluid flow patterns^25^. Both the disruption of signaling through the lacuno-canalicular network and osteocyte apoptosis have a dramatic impact in bone mechanosensation of the damaged regions and beyond, as network signal propagation is interrupted^26^. In addition, osteocytes that would be buried inside mineralized tissue now become exposed and could get hijacked by tumor cells^27^. To sum up, combining advanced multimodal 3D imaging and *ex vivo* correlative tissue characterization, we could detect, localize and characterize alterations of the ECM at the onset of small bone metastatic lesions.

Besides the metastatic lesions, we also investigated the proliferative status and localization of disseminated breast cancer cells in the bone marrow. High proliferation is a sign for an outgrowing metastasis, often stained with Ki67 proliferation marker. Miller et al. showed that the absence of Ki67 is a clear signal that a cell is in G0/G1 phase^28^. Therefore, Ki67 is an important indicator of the proliferative status of single breast cancer cells and small clusters. We found that only half of the cell clusters analyzed had one or more Ki67^+^ cell; the other half were entirely Ki67^-^. This is in agreement with previous studies describing the bone marrow as a quiescence-inducing niche^29^. In addition, breast cancer cells and small clusters were found to be located close to and within capillaries and blood vessels, with only few cells located further away. This is also in agreement with previous work showing that breast cancer cells in the bone marrow preferentially co-localize to blood vessels compared to osteoblasts or megakaryocytes^29^. The perivascular niche has been described as both activating and dormancy inducing^29, 30^, while others have proposed the endocortical niche as quiescence inducing, following the mechanisms of hematopoietic stem cells^31^. Our 3D LSFM visualization shows that eGFP^+^ bone-tropic breast cancer cells and small clusters (>5,500 of 5 different bones) are located close to the endosteal bone surface, while the 2D immunofluorescence analysis locates them in proximity with capillaries and blood vessel. Since recent work revealed that cortical bone is highly vascularized by transcortical vessels^15^, the endocortical and perivascular niche can spatially overlap and is therefore difficult to separate between the two.

Finally, we studied the altered bone (re)modeling in tumor-injected animals without detectable osteolytic lesions for up to five weeks post injection, to investigate the systemic effect of circulating breast cancer cells or established metastases in other regions. Trabecular bone in both femur and tibia showed significantly increased mineralization in volume and surface at later time points (31 days onward). The differences were more pronounced in the tibia, which is also more (re)modeled in the physiological state^20^. In addition, we found that the erosion is slightly down regulated in the tibia, in particular in the MRR. These findings show a clearly different trend compared to animals with large osteolytic disease^19^. Similar observations were reported by Chiou *et al*, who showed increased bone mineralization in animals injected with tumor conditioned media, suggesting that the increased mineralization is not caused by direct cancer cell contact; instead it could precede osteolytic disease^8^. The mechanism behind remains unclear, with arguments in favor of the premetastatic^8^ vs. anti-metastatic niche^32^.

In summary, using tissue clearing of 3D (intact) bones and LSFM we showed that breast cancer cells and small clusters home in all bone compartments, with a preference for small clusters in the bone marrow and larger clusters on the periosteum. Osteolytic lesions, on the other hand, occurred only in characteristics locations of high bone (re)modeling in the metaphysis. Based on *in vivo* microCT-based time-lapse morphometry and a new image analysis tool, we detected three types of lesions: cortical lesions on the periosteal side, lesions in the endocortical bone propagating to trabecular bone, and trabecular lesions adjacent to the growth plate. These findings support the hypothesis that dynamic bone (re)modeling is a driver for metastasis and gives breast cancer cells the ability to establish an osteolytic bone disease, as cancer cells reach every bone compartment, yet lesions only occur in regions of high bone (re)modeling in the metaphysis. Furthermore, a multiscale correlative tissue characterization of early small bone metastatic lesions showed that not only the mineral but also the collagen fibers are fully resorbed, causing severe changes in the osteocytes and the lacuno-canalicular network. The cellular analysis revealed that half of the breast cancer cells and clusters in the bone marrow are non-proliferative and located close or even within blood vessels. Further, we showed that bone (re)modeling is altered in the absence of visible osteolytic lesions, leading to higher mineralization. Our work provides novel insights of breast cancer cell homing to bone and important 3D dynamic information of bone (re)modeling in the onset of early osteolytic lesions, metastasis initiation and progression.

## Materials and Methods

### Animal model

12-week-old female BALB/c nude mice (CAnN.Cg-Foxn1nu/Crl, Charles River, Sulzfeld, Germany) were received and acclimatized in the animal facility of the Charité - Universitätsmedizin Berlin. The mice were housed 2-4 animals per cage with ad libitum access to food and water. A reference microCT scan was acquired at day 0 and mice were injected after 4 days, under ultrasound guidance (Vevo2100, FUJIFILM VisualSonics Inc., Canada), into the left ventricle of the heart (one injection per mouse) using a 27G needle (Fig. S3A). The control mice (n=10) were injected with PBS to ensure compatibility after this stressful procedure, with 3 animals being sacrificed at day 17 for better comparison with tumor animal and *ex vivo* experiments, keeping the age of the animals and the exposure to microCT radiation as similar as possible. The tumor animals (n = 11) were injected with MDA-MB-231-1833 BoM cells (5x10^5^ in 100 µL ice cold PBS). The animals received Carprofen (CP-Pharma Handelsgesellschaft mbH, Burgdorf, Germany) and Buprenorphin (CP-Pharma Handelsgesellschaft mbH, Burgdorf, Germany) as analgesic drugs while under anesthesia. During *in vivo* imaging and the intracardiac injection animals were anesthetized using isoflurane (CP-Pharma Handelsgesellschaft mbH, Burgdorf, Germany) at 1-2% with oxygen as a carrier, and the eyes were protected from drying with Pan-Ophtal gel (Dr. Winzer Pharma GmbH, Berlin, Germany). The initial average animal weight was 19.7 g ± 1.7 g for control animals and 20.3 g ± 1.9 g for tumor animals. The weight at sacrifice for the tumor animals was 19.2 g ± 2.0 g and for control animals, 19.6 g ± 2.1 g. All animal experiments were carried out according to the policies and procedures approved by local legal research animal welfare representatives (LAGeSo Berlin, G0224/18).

### Cell culture

MDA-MB-231-1883 BoM cells were provided by Dr. Joan Massagué and purchased from the Antibody and Bioresource Core Facility of the Memorial Sloan Kettering Cancer Center, USA^33^. Briefly, MDA-MB-231 (ATCC ® HTB-26™) human epithelial breast cancer cells were stably transduced with a lentivirus expressing a triple-fusion reporter, forming MDA-MB-231 TGL cells^34^. The here used subclone 1833 is a bone-tropic human cell line deriving from a metastasis formed by these MDA-MB-231 TGL cells hosted in a mouse. MDA-MB-231-1883 BoM cells were cultured in low glucose Dulbecco’s Modified Eagle’s Medium (DMEM) (Sigma-Aldrich, Taufkirchen, Germany) supplemented with 1% penicillin/streptomycin (Gibco) and 10% fetal bovine serum (FBS superior, Sigma-Aldrich, Taufkirchen, Germany). The cells were grown at 37 °C with 5% CO_2_ and regular passaging.

### *In vivo* microCT time-lapse image acquisition

Longitudinal imaging of the hind limbs of the mice was performed with a high resolution microCT (U-CTHR, MILabs, Netherlands). The X-ray tube was operated at 50 kVp source voltage and 210 mA source current. Images were acquired at a step angle of 0.375° with 2 projections per step (75 ms exposure time), over a range of 360° and with 8.5 µm voxel size. Two aluminum filters with 100 µm and 400 µm thickness were used. The scan region included the entire femur and tibia of both hind limbs and was determined on an X-ray scout view. To prevent motion artifacts and ensure reproducibility, the anesthetized mice were positioned using an animal bed with the hindlimbs restrained. The scans were reconstructed using the MILabs Reconstruction software and a projection filter (Hann) was applied in the process to limit blurring. The scanner was calibrated before every scan using the internal calibration system. The animals were scanned with the microCT for a reference scan (day 0, 12-week-old). After 17 days, the animals were scanned twice per week up until maximum 38 days (on days 17, 20, 24, 27, 31, 34 and 38) (Fig. S3A). Anesthetized mice were sacrificed by cervical dislocation. The hind limbs were harvested and fixed with 4% paraformaldehyde (PFA) in PBS over night at 4 °C and stored in PBS until further processed.

### *In vivo* microCT-based time-lapse morphometry

The analysis was performed in accordance with Birkhold et al.^21, 35^ and Young et al.^20^. To assess morphological changes, microCT scans of each time point (day 17, 20, 24, 27, 31, 34 and 38) were compared to the reference scan at day 0, which ensures full information for all time points and no loss of events by changing the reference scan. We analyzed both limbs of all animals. The regions of interest (ROIs) were set at the distal femoral metaphysis and the proximal tibial metaphysis, reaching 10% of the bone length. Each region of interest (ROI) started at the end of the non-mineralized growth plate, on the interface where the primary spongiosa of the metaphysis is found, reaching towards the diaphysis. The ROI included both primary and secondary spongiosa of the metaphysis. The entire epiphysis and other connected bones were manually segmented from the metaphysis. The threshold to remove the background was determined using the 3D Otsu-Method ^36^ (tool provided by the software Amira 2020.1, Thermo Fisher Scientific, MA, USA). A list of all used thresholds is reported in the Supplementary Table S1 – S4 for PBS-injected control animals and in Supplementary Table S5 – S8 for tumor animals.

The registration process and following evaluation was performed in accordance with Birkhold et al.^21^ and Young et al.^20^. The images were pre-processed and aligned using Amira. During the pre-processing step, the dataset was pre-cropped to the respective ROI. The epiphysis and other connected bones such as the fibula were removed by manual segmentation. Later time point images were registered onto the reference image (day 0) using a 3D rigid registration algorithm. The threshold was used to exclude background noise from the registration, while keeping the grey scale of the respective bone region data set. A hierarchical strategy was chosen for the registration to avoid local minima, starting with coarse resampling and proceeding to finer resolutions under visual control to ensure correct registration. After the registration, the images were transformed into the same coordinate system using a Lanczos interpolation, keeping the voxel size constant. The images were then re-cropped according to the corresponding ROI.

Next, the ROIs were segmented into trabecular and cortical bone using morphological operations as described by Birkhold et al.^21^ and evaluated, both using custom-made MATLAB (MATLAB 2018a; The Mathworks, Inc. Natick, MA, USA) scripts (Fig. S3B). Bone morphological changes were evaluated as previously described by Young et al^20^: MV/BV (normalized parameters are divided by the respective bone volume or bone surface of the reference scan), normalized mineralized bone surface-to-soft tissue interface (MS_ST_/BS), normalized mineralized bone surface-to-constant bone interface (MS_CB_/BS), MAR (mean thickness of formed bone in μm/day), EV/BV, normalized eroded bone surface-to-soft tissue interface (ES_ST_/BS) normalized eroded bone surface-to-constant bone interface (ES_CB_/BS), as well as MRR (mean depth of resorbed bone in μm/days). For each case, only the concerning voxels labeled as constant, mineralized or eroded voxels, were taken into consideration. The first layer of voxels was disregarded due to partial volume effects. The second most outward layer of voxels was considered as the soft tissue interface, while the constant bone interface was evaluated by expanding the constant bone surface by one voxel. The description of the surfaces did not influence the analysis of volumetric parameters, which were calculated from all bone voxels independently. A list of all parameters is provided in Supplementary Table S9.

### Eroded bone patch image analysis tool to detect and track osteolytic lesions in cortical bone

To detect and track the growth of osteolytic lesions in cortical bone with dynamic microCT-based time-lapse morphometry data, a new image analysis tool was developed that could differentiate between physiological and pathological erosion patches in cortical bone. Only eroded voxels of previously segmented cortical bone were considered. A chosen 3D neighborhood of each eroded voxel was evaluated, and the status of surrounding voxels – being eroded or not eroded – plus the voxel itself were considered leading to a sum matrix. Two different empirically determined thresholds were applied a) to analyze the volume of the entire eroded bone patch and b) to detect the eroded bone patch core (main criteria to identify a lesion) To analyze the volume of the eroded bone patch, the noise in form of scattered eroded voxels around the eroded bone patch core was filtered: To do so, the neighborhood of each eroded voxel was explored in 3 voxels in positive and negative x-, y- and z-direction. If a threshold of 70% was fulfilled, those erosion voxels were maintained and contributed to the volume of the eroded bone patch. The threshold parameters were determined empirically under visual control. To detect the eroded bone patch core of only pathological erosions, the neighborhood of each eroded voxel was explored in 9 voxels in positive and negative x- and y-direction and in 4 voxels in positive and negative z-direction. If a threshold of 92.61% was fulfilled, meaning that 92.61% of the neighborhood voxels were erosion voxels, that voxel was identified as belonging to the eroded bone patch core. The parameters were determined empirically ensuring that no lesions were detected in PBS-injected control animals. Only erosion patches containing a core were identified as pathological eroded bone patches. The volume of each eroded bone patch in mm^3^ was obtained at different time points. The location of osteolytic lesions was visualized with the eroded bone patch core shown in pink and the respective entire patch in green.

### Lesion analysis in trabecular bone

Trabecular osteolytic lesions were detected visually both in the raw data and the microCT-based time-lapse morphometry data. A region was considered as a lesion if the mineralized part of the growth plate was fully resorbed, leaving an abnormal hole in the spongy structure of the mineralized growth plate.

### Code availability

The original code as well as the modified version used in this study are available on GitHub: https://github.com/BWillieLab/Timelapse-Morphometry (original code, Birkhold et al.^21, 35^) https://github.com/CipitriaLab/Timelapse-Morphometry (modified code used in this and the previous study, Young et al.^20^) https://github.com/CipitriaLab/Timelapse-Morphometry/tree/lesion_analysis (eroded bone patch image analysis tool developed in this study)

### Tissue clearing and optical light-sheet fluorescence microscopy (LSFM)

Fixed tibiae were dehydrated in an ascending ethanol series (50, 70 and two times in 100%) for 12 h each at room temperature and cleared with ethyl cinnamate (Sigma-Aldrich, Taufkirchen, Germany) at room temperature for 24 h according to the simpleCLEAR protocol^15^ (Fig. S3C). To image cleared tibiae via LSFM, a LaVision BioTec Ultramicroscope Blaze (Miltenyi/LaVision BioTec, Bielefeld, Germany) with a supercontinuum white light laser (460 – 800 nm), 7 individual combinable excitation and emission filters covering 450 – 865 nm, an AndorNeo sCMOS Camera with a pixel size of 6.5 μm^3^, and 1.1× (NA 0.1), 4× (NA 0.35), 12× (NA 0.53) objectives, with a magnification changer ranging from 0.66× to 30× was used. For image acquisition, cleared samples were immersed in ethyl cinnamate in a quartz cuvette and excited at 470/30 nm for eGFP-excitation and 560/40 nm for tissue autofluorescence excitation. eGFP was detected using a 525/50 nm band-pass emission filter and a 620/60 nm band-pass emission filter was used for tissue autofluorescence detection. Because the excitation optics of the microscope provide a light-sheet thickness of 4 – 24 μm (adjustable NA), the Z-step size was set to 5 or 10 μm depending on the selected light-sheet NA. The optical zoom factor varied from 1.66× to 4× depending on the objectives and digital zoom factors used.

Imaris (Bitplane, Version 9.6.0, Oxford Instruments, Abingdon, UK) was used for data analysis. The bone marrow was segmented manually and the cells were analyzed using the spots tool with an initial estimated diameter of 35.5 μm that was found by manually measuring around 10 cells. n = 5 bones from five different animals were analyzed.

The clearing protocol was reversed by placing samples in a descending ethanol series (100, 70, 50%) with 1% Tween-20 for 5 h at room temperature each, followed by two steps in PBS with 1% Tween-20 (5 h at room temperature each). The samples were stored in PBS until further use.

### Resin embedding of mineralized bone tissue

Bone specimens were cold embedded at 4 °C in poly(methyl methacrylate) (PMMA) (Technovit 9100, Kulzer, Germany), following the manufacturer’s instructions. Briefly, samples were dehydrated in an ascending ethanol series (70, 80, 96 and 100%) for 48 h at 4 °C each with a final step in rhodamine solution (4.17 mg/mL−1 Rhodamine B, pure in 100% ethanol, ACROS Organics, Geel, Belgium), followed by a xylene washing, pre-infiltration and infiltration, each for 48 h at 4 °C, and final embedding in PMMA^24^.

### Sectioning using *ex vivo* microCT and controlled angle cutting

In order to expose the metastatic osteolytic lesion, we used controlled angle cutting as described by Moreno-Jimenez et al.^24^. The excess PMMA was first trimmed and the entire block scanned with an *ex vivo* microCT scanner (EasyTom micro/nano tomograph RX solutions, France) at a low resolution with 15 μm voxel size and acquisition parameters of 107 kV, 93 μA, frame rate of 6 and average frame of 5. Reconstruction of the 800 projections/scan were performed using RX Solutions X-Act software. The cutting plane to capture the metastatic osteolytic lesion was found using Data Viewer (Bruker, Kontich, Belgium). For small correction angles under 20°, a tilted cylinder was cut and the sample blocks were fixed on it for sectioning. For larger correction angles over 20°, an additional cut of the block was performed with a diamond wire saw (DWS.100, Diamond WireTec GmbH & Co.KG, Weinheim, Germany) and a second microCT scan was performed, repeating the same procedure as above. The histological sections were taken at a thickness of 6 μm with a microtome (Leica RM2255 Microtome, Leica Biosystems, Wetzlar, Germany).

### Haematoxylin and eosin staining of PMMA sections

The PMMA embedded sections were stained using a standard protocol of haematoxylin and eosin (H&E) (H&E Rapid kit, Clin-Tech, Surrey, UK). In short, samples were rehydrated in a descending ethanol series, followed by staining in Carazzi’s double-strength haematoxylin. After another wash in tap water, the slides were stained in eosin, rinsed in tap water, dehydrated in an ascending ethanol series (70, 80, 96, 100% twice for 2 min each), incubated in xylene twice for 2 min and mounted (VITROCLUD, deltalab, Barcelona, Spain). The stained sections were imaged with a Keyence Digital Microscope (VKX-5550E, Keyence, Germany). Mineralized tissue and nuclei are shown in dark purple, while soft tissue is shown in light purple.

### Confocal laser scanning microscopy (CLSM) of PMMA block surface

After sectioning, the exposed PMMA block surface was polished (PM5, Logitech, Glasgow, Scotland) with abrasive paper with grading 4000, and diamond spray with grading of 5 μm, 3 μm and 1 μm, under constant visual control, ensuring that only minimal tissue got removed. The block surface of the rhodamine-stained samples was imaged using CLSM (Leica TCS SP8 DLS, Multiphoton, Leica Microsystems CMS GmbH, Wetzlar, Germany). The samples were imaged with a HC PL APO CS2 40×/1.30 oil objective (NA 1.3) at a magnification of 40×. Argon laser light with λ_excitation_ = 514 nm / λ_emission_ = 550 – 650 nm was used with a laser intensity of 10%. Images were acquired with a resolution of 1024 x 1024 pixels (pixel size of 284 nm) at a scan speed of 400 Hz. Over 100 images for each sample with a thickness of 347 nm, spanning a changing thickness of around 50 μm depth from the block surface were imaged. To compensate for the loss of signal with increasing depth, the photomultiplier gain was continuously increased. The images were stitched and visualized with the LAS X software (Version 3.5.7.23225, Leica Microsystems CMS GmbH, Wetzlar, Germany). For each animal, a region containing an osteolytic lesion, either in the cortical or trabecular bone, and a lesion-free region in the same bone at a comparable location were imaged. n = 5 bones from five different animals were analyzed.

### Second-harmonic generation (SHG) correlative imaging of PMMA block surface

SHG imaging was performed on the exact same ROI with the same instrument and objective. A pulsed infrared laser (λ_excitation_ = 910 nm/ λ_emission_ = 450 – 460 nm) with a power of 0.6 W and an intensity of 10% was used. Images were acquired with a resolution of 1024 x 1024 pixels (pixel size of 284 nm) at a scan speed of 400 Hz. Around 20 images with a thickness of 2.8 μm were acquired, keeping the same total depth as the previous rhodamine-stained fluorescence imaging. For each animal a region containing an osteolytic lesion, either in the cortical or trabecular bone, and a lesion-free region at a comparable location were imaged. The images were stitched and visualized with the LAS X software (Version 3.5.7.23225, Leica Microsystems CMS GmbH, Wetzlar, Germany). n = 5 bones from five different animals were analyzed.

### Backscattered electron (BSE) correlative imaging of PMMA block surface

BSE imaging was performed on the exact same ROI with an environmental scanning electron microscope (Quattro ESEM, Thermo Fisher Scientific, Waltham, US). The following settings were used: 15 kV, low vacuum conditions (100 Pa), working distance of approximately 8 mm, spot size of 3 and 500× magnification. The images were manually stitched using Adobe Photoshop 2022. n = 5 bones from five animals were analyzed.

### Cryo-embedding and -sectioning of mineralized bone tissue

Bones of one limb of each mouse without a detectable osteolytic lesion were freeze-embedded following the method of the SECTION-LAB Co. Ltd. (Hiroshima, Japan). Briefly, the samples were dehydrated in an ascending sucrose solution (10, 20 and 30% in distilled water) for 24 h each at 4 °C. Next, a metal mold was placed in isopropanol, cooled with liquid nitrogen and filled with embedding medium (SCEM; SECTION-LAB Co. Ltd., Hiroshima, Japan). The bone was placed in the middle and covered with embedding medium, avoiding direct contact with the cooled isopropanol. Samples were stored at −80 °C until further use. Cryosections with a thickness of 20 μm were cut following the Kawamoto method using a cryostat (Leica CM3060S, Leica Microsystems CMS GmbH, Wetzlar, Germany). The sections were collected using Kawamoto films (type 2C(9), SECTION-LAB Co. Ltd., Hiroshima, Japan), then attached to a microscopic slide and stored at -20 °C until further use.

### Immunofluorescence staining and imaging of cryosections

For IF staining, the slides were blocked with blocking buffer (1% bovine serum albumin (BSA), Sigma-Aldrich, Taufkirchen, Germany, 0.1% Tween20 Carl Roth, Karlsruhe, Germany, 0.1% dimethyl sulfoxide (DMSO), Merck KGaA, Darmstadt, Germany in PBS) for 1 h. Primary and secondary antibodies were diluted in blocking buffer and incubated for at least 4 h at room temperature or overnight at 4 °C, with in between washing steps (three times, 15 min each) in washing buffer (0.1% Tween20, 0.1% DMSO in PBS). Final washing steps were conducted in washing buffer (twice for 15 min) and in distilled water (once for 15 min). Slides were then mounted with Dako fluorescence mounting media (S302380-2, Agilent Technologies, Santa Clara, US). The primary antibodies and respective concentrations used in this study are the following: 4’,6-diamidino-2-phenylindole (DAPI) (1:500, Roche), anti-Ki67 (1:100, 14-5698-82, eBioscience™) and anti-CD31 (1:200, 102516, Biolegend). As secondary antibody Alexa Fluor 555 (1:1000, 4417S, Cell Signaling Technology) was used. Samples were imaged using CLSM (Leica TCS SP8 DLS, Multiphoton, Leica Microsystems CMS GmbH, Wetzlar, Germany).

Three different objectives were used: HC APO L 10x/0.30 water objective (NA 0.3) at a magnification of 10×, HC PL APO CS2 40×/1.30 oil objective (NA 1.3) at a magnification of 40×, HC PL APO CS2 63x/1.40 oil objective (NA 1.4) at a magnification of 63×. For imaging the DAPI, diode 405 laser light with λ_excitation_ = 405nm / λ_emission_ = 410 – 483nm was used. For imaging the eGFP, argon laser light with λ_excitation_ = 488nm / λ_emission_ = 493 – 556nm was used. For imaging the Alexa Fluor 555, DPSS 561 laser light with λ_excitation_ = 561nm / λ_emission_ = 566 – 756nm laser light was used. For imaging the Alexa Fluor 647, HeNe 633 laser light with λ_excitation_ = 633nm / λ_emission_ = 638 – 755nm was used. Laser intensity and photomultiplier gain were kept constant: DAPI (intensity 1%, HyD detector, gain 30), eGFP (intensity 20%, Hyd SMD 2 detector, gain 2900), Alexa Fluor 555 (intensity 1%, PMT detector, gain 650) and Alexa Fluor 647 (intensity 3%, HyD detector, gain 200). Images were acquired with a resolution of 1024 x 1024 pixels (pixel size of 284 nm) at a scan speed of 400 Hz (200 Hz for 63× magnification). A varying number of images with a thickness of 4 μm (2 μm for 63× magnification) were made over a depth of around 20 μm. For 10× and 40× magnification, tile scans were acquired depending on the size of the sample.

Images were analyzed and visualized with Imaris (Imaris Viewer 9.8.0, Oxford Instruments, Abingdon, United Kingdom). 40× magnification images were used for quantification and the thresholds were applied manually under visual control. For Ki67 analysis, cells/clusters were manually segmented using the surface tool on each layer. Nuclear Ki67 signal was overlapped using the DAPI signal and counted automatically. For distance analysis with CD31, cancer cells were estimated on one layer using the surface tool. CD31 was segmented using the surface tool, distance mapping and overlapping were calculated automatically. n = 3 (Ki67) and n = 4 (CD31) bones from four animals were analyzed.

### Statistical analysis

In the analysis of altered bone (re)modeling in absence of detectable bone lesions, only bones harvested from animals without detectable osteolytic lesions in the hind limbs were included. All bones harvested from animals with confirmed lesions were excluded from this analysis. Both limbs of all animals were considered, resulting in seven control animals with n = 14 bones and 5 tumor animals with n = 10 bones, for femur and tibia respectively. The plots show the mean and standard deviation. Two sample t-tests were used for statistical comparison between different groups. Significant differences are presented as * p ≤ 0.05.

## Acknowledgements

This work was funded by the Deutsche Forschungsgemeinschaft (DFG) Emmy Noether grant (CI 203/2-1 to A. C., S. A. E. Y., and D. S. G.). A. C. also thanks the funding from IKERBASQUE Basque Foundation for Science and from the Spanish Ministry of Science and Innovation (PID2021-123013OB-I00) (Ministerio de Ciencia e Innovación, la Agencia y del Fondo Europeo de Desarrollo Regional, Proyecto PID2021-123013OB-I00 financiado por MCIN/AEI/10.13039/501100011033/FEDER.UE). A.-D. H. thanks the International Max Planck Research School (IMPRS) on Multiscale Bio Systems for financial support. The FRQS Programme de bourse de chercheur provided support to B.W. We thank the lab of Dr. Joan Massagué at the Memorial Sloan Kettering Cancer Center, USA, for providing the MDA-MB-231-1883 BoM cells. We thank Jeannette Steffen for technical assistance with sectioning and staining. The authors would like to acknowledge Peter Fratzl, Sadra Bakhshandeh and Hubert Taïeb for scientific discussion.

## Author contribution

Study design: AC, SAEY

Experimental work: SAEY, DSG, VQ, AE, AC

Data analysis: SAEY, ADH, AG, AC

Visualization: SAEY, ADH, AC

Data interpretation: SAEY, ADH, MR, GND, BMW, AG, AC

Project supervision: AC

Drafting manuscript: SAEY, AC

Revising manuscript content and approving final version: all authors

## Data and materials availability

The MATLAB codes used are freely available in GitHub and the links are provided in the Methods section. The data supporting the findings can be made available upon reasonable request.

## References

1. Torre, L. A. et al. Global cancer statistics, 2012. CA. Cancer J. Clin. 65, 87–108 (2015).

2. Coleman, R. E. Clinical Features of Metastatic Bone Disease and Risk of Skeletal Morbidity. Clin. Cancer Res. 12, 6243s–6249s (2006).

3. Weilbaecher, K. N., Guise, T. A. & McCauley, L. K. Cancer to bone: a fatal attraction. Nat. Rev. Cancer 11, 411–425 (2011).

4. Harbeck, N. et al. Breast cancer. Nat. Rev. Dis. Prim. 5, 66 (2019).

5. Coleman, R. E. et al. Bone metastases. Nat. Rev. Dis. Prim. 6, 83 (2020).

6. Phan, T. G. & Croucher, P. I. The dormant cancer cell life cycle. Nat. Rev. Cancer 20, 398–411 (2020).

7. Croucher, P. I., McDonald, M. M. & Martin, T. J. Bone metastasis: the importance of the neighbourhood. Nat. Rev. Cancer 16, 373–386 (2016).

8. Chiou, A. E. et al. Breast cancer–secreted factors perturb murine bone growth in regions prone to metastasis. Sci. Adv. 7, 1–14 (2021).

9. Crist, S. B. & Ghajar, C. M. When a House Is Not a Home: A Survey of Antimetastatic Niches and Potential Mechanisms of Disseminated Tumor Cell Suppression. Annu. Rev. Pathol. Mech. Dis. 16, 409–432 (2021).

10. Price, T. T. et al. Dormant breast cancer micrometastases reside in specific bone marrow niches that regulate their transit to and from bone. Sci. Transl. Med. 8, 340ra73–340ra73 (2016).

11. He, F. et al. Multiscale characterization of the mineral phase at skeletal sites of breast cancer metastasis. Proc. Natl. Acad. Sci. 114, 10542–10547 (2017).

12. Di Martino, J. S. et al. A tumor-derived type III collagen-rich ECM niche regulates tumor cell dormancy. *Nat*. Cancer 2021 1–18 (2021). doi:10.1038/s43018-021-00291-9

13. Fonta, C. M. et al. Fibronectin fibers are highly tensed in healthy organs in contrast to tumors and virus-infected lymph nodes. Matrix Biol. Plus 8, 100046 (2020).

14. Grüneboom, A. et al. Next-generation imaging of the skeletal system and its blood supply. Nat. Rev. Rheumatol. 15, 533–549 (2019).

15. Grüneboom, A. et al. A network of trans-cortical capillaries as mainstay for blood circulation in long bones. Nat. Metab. 1, 236–250 (2019).

16. Winkler, J., Abisoye-Ogunniyan, A., Metcalf, K. J. & Werb, Z. Concepts of extracellular matrix remodelling in tumour progression and metastasis. Nat. Commun. 11, 5120 (2020).

17. Fratzl, P. & Weinkamer, R. Nature’s hierarchical materials. Prog. Mater. Sci. 52, 1263– 1334 (2007).

18. Wang, N. et al. The frequency of osteolytic bone metastasis is determined by conditions of the soil, not the number of seeds; evidence from in vivo models of breast and prostate cancer. J. Exp. Clin. Cancer Res. 34, 124 (2015).

19. Johnson, L. C. et al. Longitudinal live animal micro-CT allows for quantitative analysis of tumor-induced bone destruction. Bone 48, 141–151 (2011).

20. Young, S. A. E. et al. In vivo microCT-based time-lapse morphometry reveals anatomical site-specific differences in bone (re)modeling serving as baseline parameters to detect early pathological events. Bone 161, 116432 (2022).

21. Birkhold, A. I. et al. Mineralizing surface is the main target of mechanical stimulation independent of age: 3D dynamic in vivo morphometry. Bone 66, 15–25 (2014).

22. Aceto, N. et al. Circulating Tumor Cell Clusters Are Oligoclonal Precursors of Breast Cancer Metastasis. Cell 158, 1110–1122 (2014).

23. Campbell, G. M. et al. Tracking the Progression of Osteolytic and Osteosclerotic Lesions in Mice Using Serial In Vivo μCT: Applications to the Assessment of Bisphosphonate Treatment Efficacy. J. Bone Miner. Res. 33, 410–418 (2018).

24. Moreno-Jiménez, I., Garske, D. S., Lahr, C. A., Hutmacher, D. W. & Cipitria, A. Targeted 2D histology and ultrastructural bone analysis based on 3D microCT anatomical locations. MethodsX 8, 101480 (2021).

25. van Tol, A. F. et al. Network architecture strongly influences the fluid flow pattern through the lacunocanalicular network in human osteons. Biomech. Model. Mechanobiol. (2019). doi:10.1007/s10237-019-01250-1

26. Weinkamer, R., Kollmannsberger, P. & Fratzl, P. Towards a Connectomic Description of the Osteocyte Lacunocanalicular Network in Bone. Curr. Osteoporos. Rep. 17, 186–194 (2019).

27. Hemmatian, H. et al. Reorganization of the osteocyte lacuno-canalicular network characteristics in tumor sites of an immunocompetent murine model of osteotropic cancers. Bone 116074 (2021). doi:10.1016/j.bone.2021.116074

28. Miller, I. et al. Ki67 is a Graded Rather than a Binary Marker of Proliferation versus Quiescence. Cell Rep. 24, 1105–1112.e5 (2018).

29. Carlson, P. et al. Targeting the perivascular niche sensitizes disseminated tumour cells to chemotherapy. Nat. Cell Biol. 21, 238–250 (2019).

30. Ghajar, C. M. et al. The perivascular niche regulates breast tumour dormancy. Nat. Cell Biol. 15, 807–817 (2013).

31. Dhawan, A. et al. Breast cancer cells compete with hematopoietic stem and progenitor cells for intercellular adhesion molecule 1-mediated binding to the bone marrow microenvironment. Carcinogenesis 37, 759–767 (2016).

32. Kolb, A. D., Shupp, A. B., Mukhopadhyay, D., Marini, F. C. & Bussard, K. M. Osteoblasts are “educated” by crosstalk with metastatic breast cancer cells in the bone tumor microenvironment. Breast Cancer Res. 21, 31 (2019).

33. Minn, A. J. et al. Genes that mediate breast cancer metastasis to lung. Nature 436, 518– 524 (2005).

34. Ponomarev, V. et al. A novel triple-modality reporter gene for whole-body fluorescent, bioluminescent, and nuclear noninvasive imaging. Eur. J. Nucl. Med. Mol. Imaging 31, 740–751 (2004).

35. Birkhold, A. I. et al. The influence of age on adaptive bone formation and bone resorption. Biomaterials 35, 9290–9301 (2014).

36. Otsu, N. A Threshold Selection Method from Gray-Level Histograms. IEEE Trans. Syst. Man. Cybern. 9, 62–66 (1979).

