## Supplementary information for "From breast cancer cell homing to the onset of early bone metastasis: dynamic bone (re)modeling as a driver of metastasis"

Phone number: +49 331 567-9452

**This PDF file includes:**

Figs. S1 to S3

Tables S1 to S9

Captions for Movies S1 to S4

**Other Supplementary Materials for this manuscript include the following:**

Movies S1 to S4

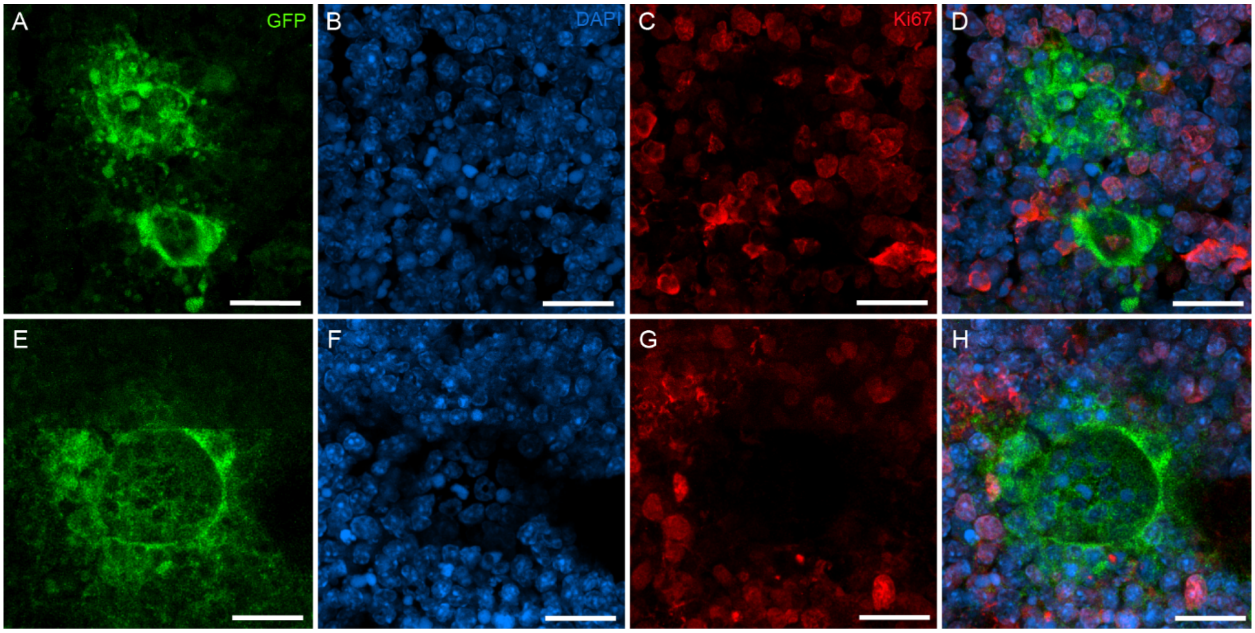


**Fig. S1: Different proliferation status of cancer cell clusters in the bone marrow.** (**A**) $eGFP^+$ cancer cell cluster in the bone marrow (green), (**B**) with corresponding nuclear DAPI signal (blue) surrounded by other bone marrow cells and (**C**) corresponding nuclear $Ki67^+$ signal (red). (**D**) Overlap of all three signals showing a $Ki67^+$ cancer cell cluster. (**E**) $eGFP^+$ cancer cell cluster in the bone marrow (green), (**F**) with corresponding nuclear DAPI signal (blue) surrounded by other bone marrow cells and (**G**) corresponding nuclear Ki67 signal (red). (**H**) Overlap of all three signals showing a $Ki67^-$ cancer cell cluster. (Scale bar corresponds to 20 μm.)

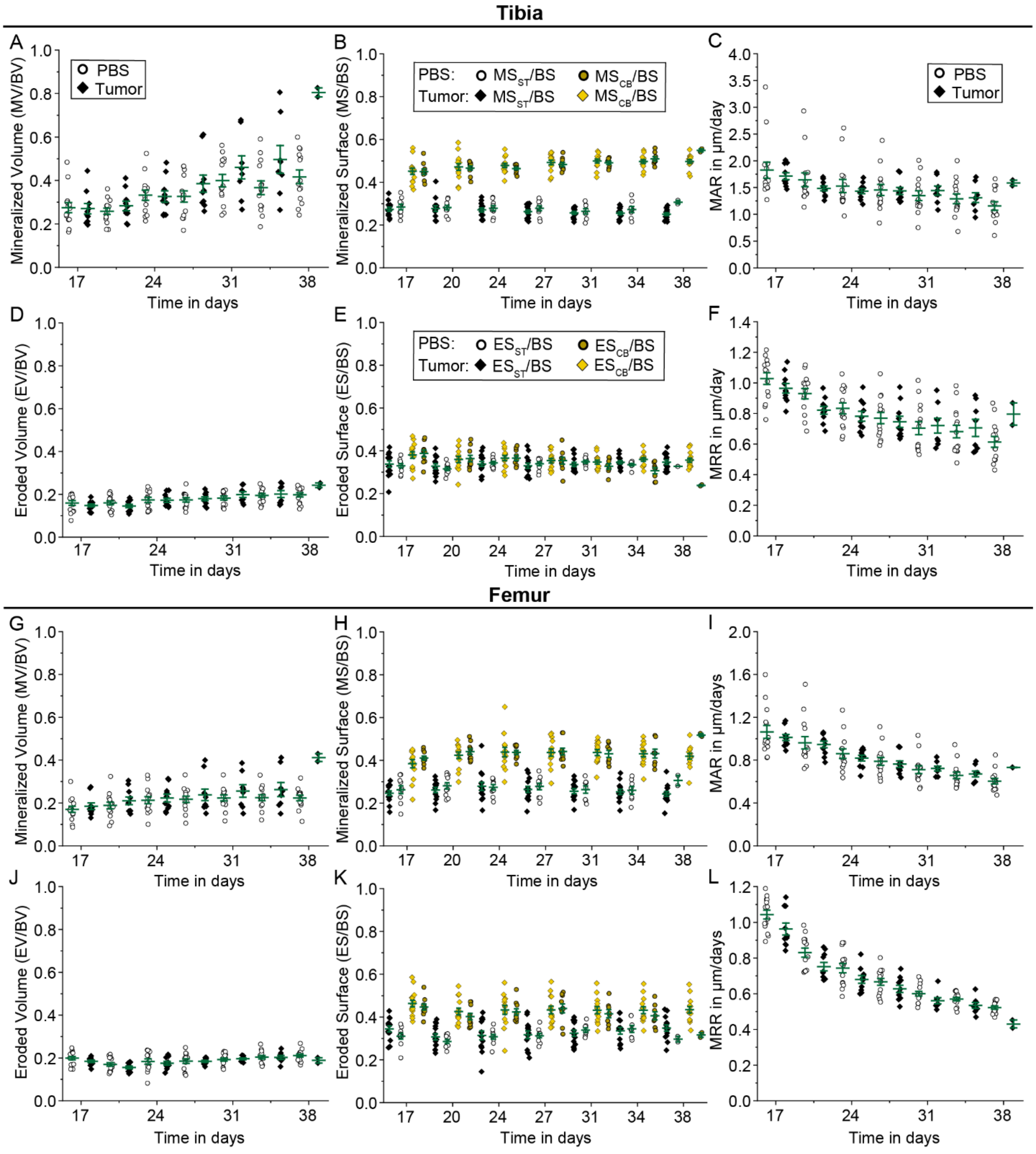


**Fig. S2: Altered cortical bone (re)modeling in absence of detectable bone lesions.** Results in cortical bone of tibial proximal metaphysis for tumor animals and PBS-injected controls are shown for mineralization with (**A**) MV/BV, (B) MS/BS and (**C**) MAR, as well as erosion with

(**D**) EV/BV, (**E**) ES/BS and (**F**) MRR. Analogous analysis in cortical bone of the femoral distal metaphysis are shown for mineralization with (**G**) MV/BV, (**H**) MS/BS and (**I**) MAR, as well as erosion with (**J**) EV/BV, (**K**) ES/BS and (**L**) MRR. (All plots show mean and standard deviation, with statistically significant values presented as * $p \geq 0.05$. PBS group: n = 14 bones from seven animals, Tumor group: n = 10 bones from five animals, for tibia and femur respectively.)

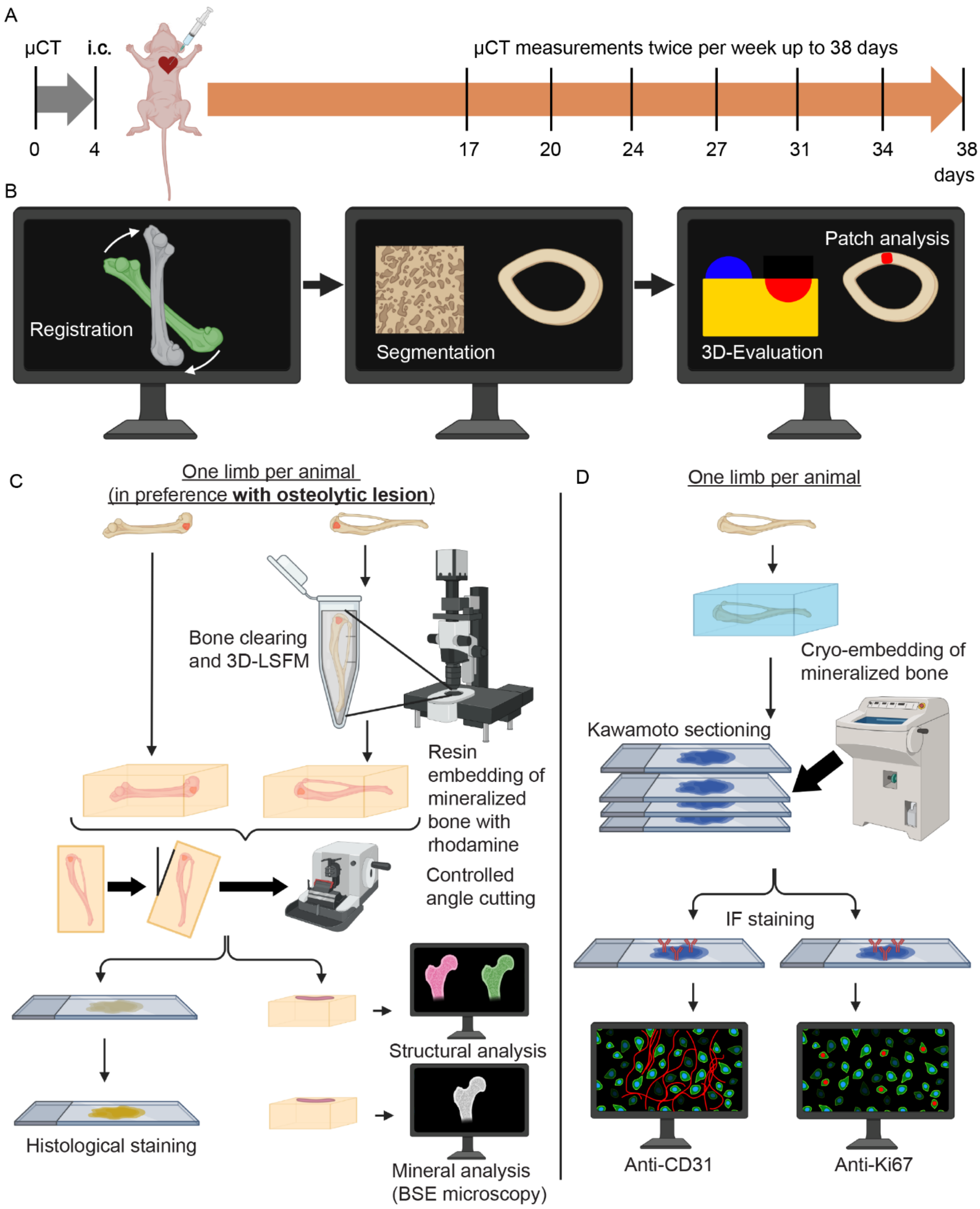


**Fig. S3: *In vivo* and *ex vivo* workflow used for multiscale correlative tissue characterization.**

(**A**) Schematic of the tumor cell injection and monitoring with *in vivo* time-lapse microCT twice per week for up to 38 days. (**B**) Depiction of the workflow for dynamic microCT-based time-lapse morphometry including registration, segmentation, evaluation and eroded bone patch analysis. (**C**) *Ex vivo* workflow for one limb per animal, preferably with an osteolytic lesion, including 3D-LSFM for tibiae, rhodamine staining and resin embedding of mineralized bones, controlled angle cutting, histological staining as well as structural and mineral analysis. (**D**) *Ex vivo* workflow for the other limb of each animal, including cryo-embedding of mineralized bones, Kawamoto sectioning and immunofluorescence (IF) staining against CD31 and Ki67 (created with BioRender.com).

**Table S1:** 3D-Otsu defined thresholds used for the left tibia ROI for each PBS control animal and time point

| **Animal / Day** | **0** | **17** | **20** | **24** | **27** | **31** | **34** | **38** |
|---|---|---|---|---|---|---|---|---|
| **1** | 1958 | 1969 | 1787 | 2126 | 2101 | 2238 | 2153 | 2245 |
| **2** | 1986 | 2131 | 2178 | 2138 | 2260 | 2219 | 2332 | 2310 |
| **3** | 2067 | 2145 | 2158 | 2173 | 2179 | 2298 | 2407 | 2495 |
| **4** | 2020 | 2081 | 2154 | 2040 | 2251 | 2271 | 2351 | 2427 |
| **5** | 1797 | 1952 | 1971 | 2058 | 1933 | 2077 | 2231 | 2287 |
| **6** | 1942 | 2062 | 2108 | 2060 | 2166 | 2184 | 2222 | 2335 |
| **7** | 1843 | 2018 | 1990 | 2054 | 2058 | 2089 | 2158 | 2245 |

**Table S2:** 3D-Otsu defined thresholds used for the right tibia ROI for each PBS control animal and time point

| Animal / Day | 0 | 17 | 20 | 24 | 27 | 31 | 34 | 38 |
|---|---|---|---|---|---|---|---|---|
| **1** | 1871 | 2071 | 2103 | 2173 | 2109 | 2269 | 2206 | 2247 |
| **2** | 1921 | 2101 | 1986 | 2199 | 2266 | 2195 | 2201 | 2241 |
| **3** | 2165 | 2072 | 2242 | 2275 | 2278 | 2327 | 2430 | 2449 |
| **4** | 2009 | 2142 | 2159 | 2230 | 2302 | 2319 | 2372 | 2410 |
| **5** | 1917 | 2008 | 1998 | 2066 | 2039 | 2125 | 2227 | 2300 |
| **6** | 1959 | 2152 | 2134 | 1989 | 2198 | 2218 | 2262 | 2337 |
| **7** | 1816 | 2070 | 2064 | 2072 | 2096 | 2127 | 2231 | 2213 |

**Table S3**: 3D-Otsu defined thresholds used for the left femur ROI for each PBS control animal and time point

| Animal / Day | 0 | 17 | 20 | 24 | 27 | 31 | 34 | 38 |
|---|---|---|---|---|---|---|---|---|
| **1** | 2276 | 2216 | 2196 | 2351 | 2202 | 2253 | 2177 | 2253 |
| **2** | 2101 | 2188 | 2205 | 2265 | 2270 | 2191 | 2307 | 2332 |
| **3** | 2080 | 2212 | 2261 | 2326 | 2342 | 2403 | 2407 | 2482 |
| **4** | 2076 | 2183 | 2225 | 2073 | 2311 | 2292 | 2338 | 2432 |
| **5** | 1957 | 2081 | 2146 | 2218 | 2205 | 2246 | 2330 | 2399 |
| **6** | 1931 | 2116 | 2134 | 2110 | 2227 | 2257 | 2300 | 2334 |
| **7** | 1935 | 2122 | 2156 | 2186 | 2255 | 2244 | 2325 | 2317 |

**Table S4:** 3D-Otsu defined thresholds used for the right femur ROI for each PBS control animal and time point

| Animal / Day | 0 | 17 | 20 | 24 | 27 | 31 | 34 | 38 |
|---|---|---|---|---|---|---|---|---|
| **1** | 2099 | 2185 | 2207 | 2274 | 2241 | 2438 | 2328 | 2353 |
| **2** | 2016 | 2148 | 2060 | 2310 | 2290 | 2237 | 2358 | 2308 |
| **3** | 2139 | 2217 | 2287 | 2408 | 2378 | 2438 | 2474 | 2499 |
| **4** | 2075 | 2233 | 2334 | 2273 | 2369 | 2305 | 2394 | 2382 |
| **5** | 2009 | 2031 | 2121 | 2235 | 2272 | 2196 | 2259 | 2293 |
| **6** | 1976 | 2152 | 2213 | 2028 | 2237 | 2257 | 2293 | 2322 |
| **7** | 1896 | 2110 | 2168 | 2194 | 2274 | 2294 | 2287 | 2342 |

**Table S5:** 3D-Otsu defined thresholds used for the left tibia ROI for each tumor animal and time point

| Animal / Day | 0 | 17 | 20 | 24 | 27 | 31 | 34 | 38 |
|---|---|---|---|---|---|---|---|---|
| **1** | 2023 | 2091 | 1991 | 2139 | 2177 | 2188 | 2149 | |
| **2** | 2057 | 2205 | 2121 | 2280 | 2369 | | | |
| **3** | 2038 | 2161 | 2187 | 2148 | 2223 | | | |
| **4** | 2082 | 2075 | 2179 | | | | | |
| **5** | 2030 | 2176 | | | | | | |
| **6** | 1834 | 2053 | | | | | | |
| **7** | 2027 | 2035 | 2097 | 2153 | 2204 | 2183 | 2196 | |
| **8** | 1918 | 1926 | 1943 | 2010 | 2036 | 2078 | 2149 | |

| | | | | | | | | |
|---|---|---|---|---|---|---|---|---|
| **9** | 1953 | 2036 | 2134 | 2183 | | | | |
| **10** | 2104 | 2113 | | | | | | |
| **11** | 1854 | 1927 | 1968 | 2037 | 1993 | 2103 | 2139 | 2170 |

**Table S6:** 3D-Otsu defined thresholds used for the right tibia ROI for each tumor animal and time point

| **Animal / Day** | **0** | **17** | **20** | **24** | **27** | **31** | **34** | **38** |
|---|---|---|---|---|---|---|---|---|
| **1** | 2089 | 2127 | 2089 | 2237 | 2246 | 2211 | 2297 | |
| **2** | 2088 | 2138 | 2131 | 2260 | 2308 | | | |
| **3** | 2071 | 2195 | 2234 | 2188 | 2121 | | | |
| **4** | 2069 | 2032 | 2200 | | | | | |
| **5** | 1993 | 1866 | | | | | | |
| **6** | 1997 | 2065 | | | | | | |
| **7** | 2018 | 2087 | 2068 | 2042 | 2196 | 2156 | 2207 | |
| **8** | 1932 | 1911 | 2005 | 2084 | 2054 | 2100 | 2131 | |
| **9** | 1922 | 2066 | 2146 | 2157 | | | | |
| **10** | 2139 | 2182 | | | | | | |
| **11** | 1917 | 1872 | 2024 | 2082 | 2075 | 2177 | 2177 | 2199 |

**Table S7:** 3D-Otsu defined thresholds used for the left femur ROI for each tumor animal and time point

| **Animal / Day** | **0** | **17** | **20** | **24** | **27** | **31** | **34** | **38** |
|---|---|---|---|---|---|---|---|---|

| | | | | | | | | |
|---|---|---|---|---|---|---|---|---|
| **1** | 2109 | 2162 | 2160 | 2255 | 2277 | 2251 | 2289 | |
| **2** | 2175 | 2346 | 2279 | 2352 | 2433 | | | |
| **3** | 2106 | 2246 | 2286 | 2304 | 2356 | | | |
| **4** | 2029 | 2196 | 2279 | | | | | |
| **5** | 2052 | 2243 | | | | | | |
| **6** | 1970 | 2201 | | | | | | |
| **7** | 2048 | 2146 | 2263 | 2259 | 2304 | 2290 | 2311 | |
| **8** | 2003 | 2043 | 2135 | 2225 | 2208 | 2222 | 2329 | |
| **9** | 2113 | 2275 | 2221 | 2290 | | | | |
| **10** | 2228 | 2233 | | | | | | |
| **11** | 1973 | 2088 | 2169 | 2233 | 2152 | 2243 | 2284 | 2299 |

**Table S8:** 3D-Otsu defined thresholds used for the right femur ROI for each tumor animal and time point

| **Animal / Day** | **0** | **17** | **20** | **24** | **27** | **31** | **34** | **38** |
|---|---|---|---|---|---|---|---|---|
| **1** | 2083 | 2215 | 2177 | 2297 | 2361 | 2338 | 2392 | |
| **2** | 2183 | 2341 | 2298 | 2375 | 2401 | | | |
| **3** | 2137 | 2203 | 2369 | 2318 | 2355 | | | |
| **4** | 2032 | 2241 | 2298 | | | | | |
| **5** | 2017 | 2066 | | | | | | |
| **6** | 1985 | 2174 | | | | | | |
| **7** | 2036 | 2121 | 2217 | 2240 | 2281 | 2230 | 2294 | |

| | | | | | | | | |
|---|---|---|---|---|---|---|---|---|
| **8** | 1958 | 2056 | 2179 | 2147 | 2190 | 2274 | 2350 | |
| **9** | 1997 | 2198 | 2188 | 2225 | | | | |
| **10** | 2172 | 2300 | | | | | | |
| **11** | 1930 | 2047 | 2107 | 2161 | 2139 | 2260 | 2259 | 2279 |

**Table S9:** List of abbreviations used in the microCT-based time-lapse morphometry analysis with definition and unit

| **Abbreviation** | **Definition** | **Unit** |
|---|---|---|
| MV/BV | Normalized mineralized bone volume | - |
| EV/BV | Normalized eroded bone volume | - |
| MS/BS | Normalized mineralized bone surface | - |
| $MS_{ST}$/BS | Normalized mineralized bone surface-to-soft tissue interface | - |
| $MS_{CB}$/BS | Normalized mineralized bone surface-to-constant bone interface | - |
| ES/BS | Normalized eroded bone surface | - |
| $ES_{ST}$/BS | Normalized eroded bone surface-to-soft tissue interface | - |
| $ES_{CB}$/BS | Normalized eroded bone surface-to-constant bone interface | - |
| MAR | Mineral apposition rate | µm/days |
| MRR | Mineral resorption rate | µm/days |

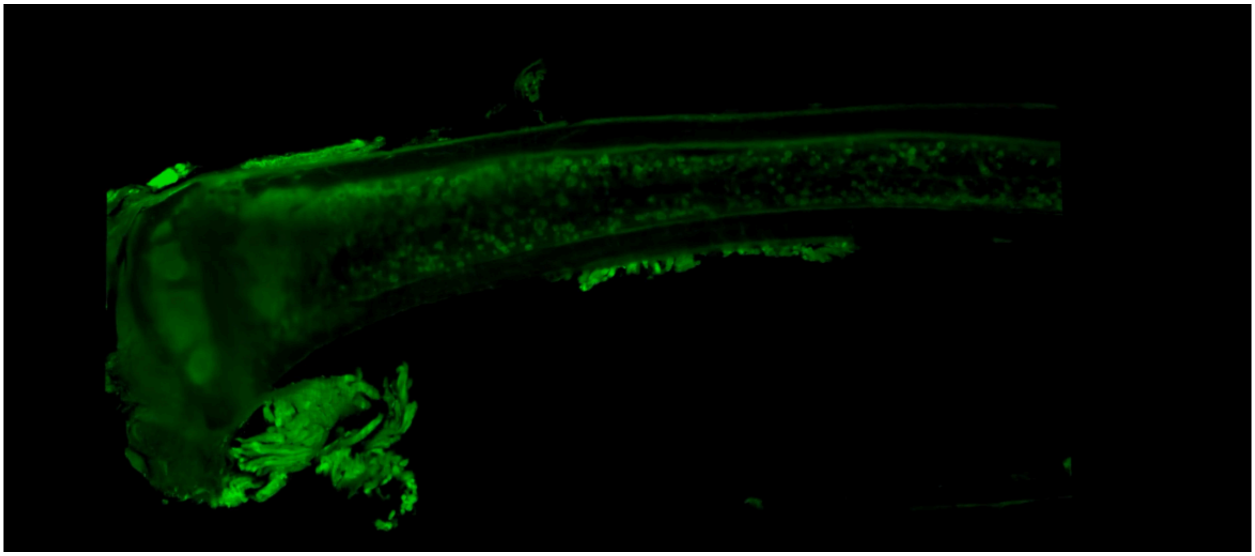

**Supplementary Movie 1**: 3D Detection of breast cancer cells in a 3D (intact) mouse tibia with light sheet fluorescence microscopy.

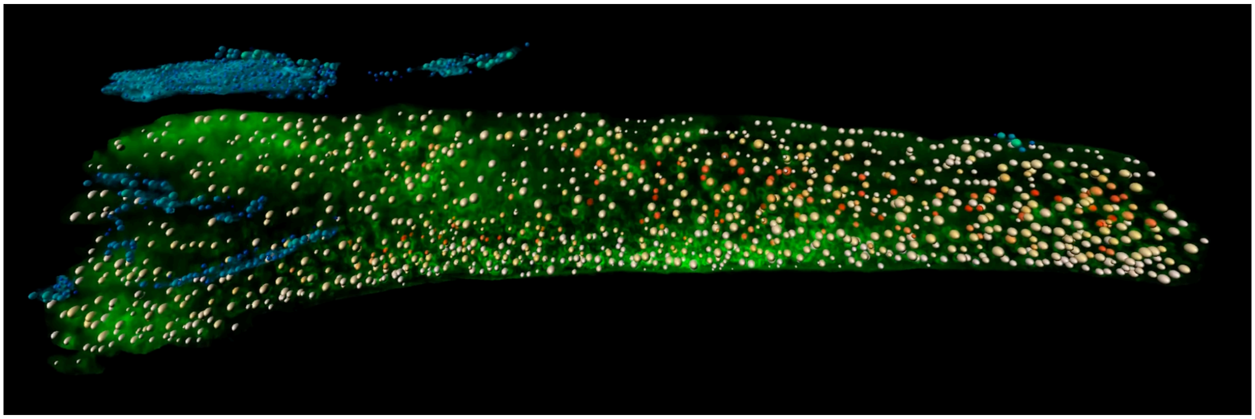

**Supplementary Movie 2**: Quantification of the size and spatial distribution of breast cancer cells and small clusters in the bone marrow of a 3D (intact) mouse tibia.

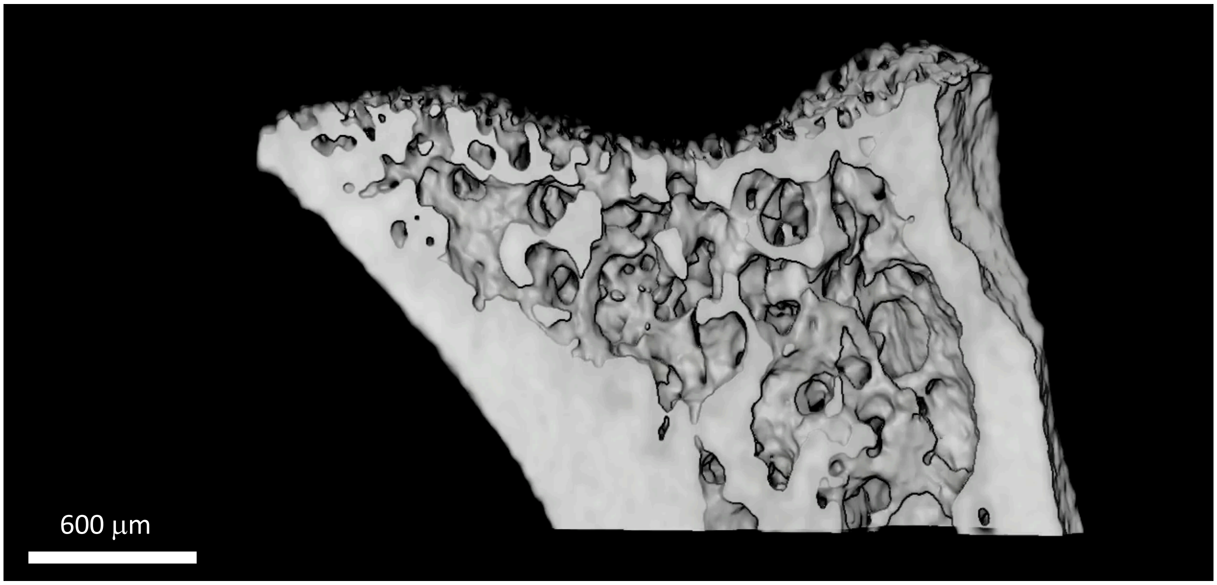


**Supplementary Movie 3**: Early bone osteolytic lesion in mouse tibia at day 27 visualized with *in vivo* microCT.

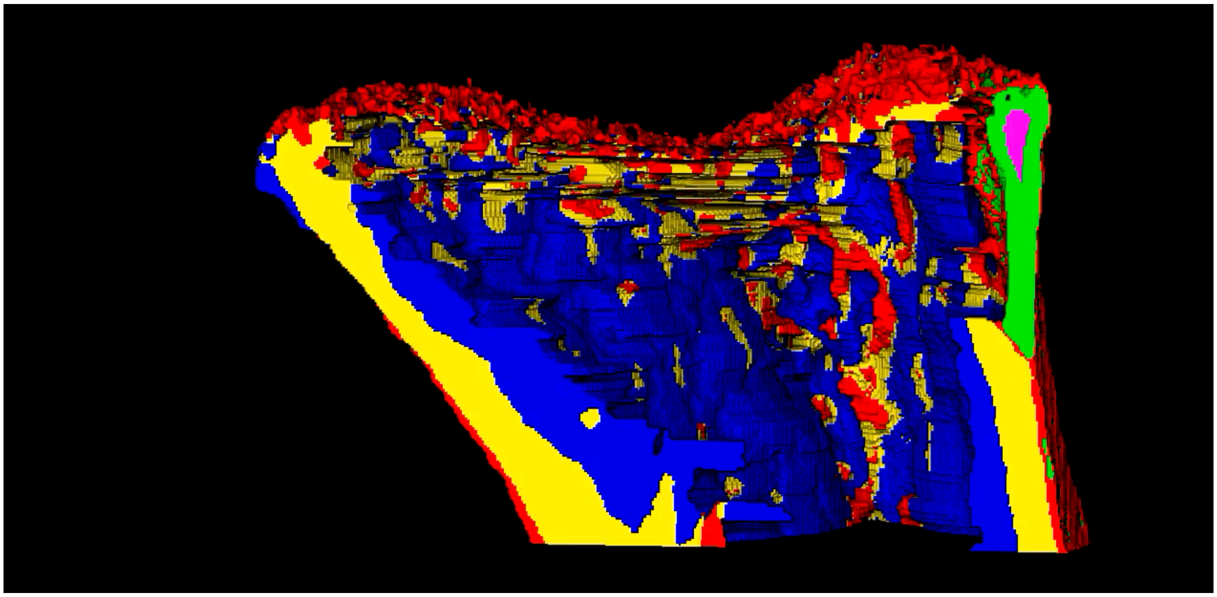

**Supplementary Movie 4**: Detection and tracking of early bone osteolytic lesions in mouse tibia with the newly developed image-based analysis tool, showing the eroded bone patch in green and the eroded bone patch core in pink.
